# Mapping the human hematopoietic stem and progenitor cell hierarchy through integrated single-cell proteomics and transcriptomics

**DOI:** 10.1101/2024.07.05.602277

**Authors:** Benjamin Furtwängler, Nil Üresin, Sabrina Richter, Mikkel Bruhn Schuster, Despoina Barmpouri, Henrietta Holze, Anne Wenzel, Kirsten Grønbæk, Kim Theilgaard-Mönch, Fabian J. Theis, Erwin M. Schoof, Bo T Porse

## Abstract

Single-cell transcriptomics (scRNA-seq) has enabled the characterization of cell state heterogeneity and recapitulation of differentiation trajectories. However, since proteins are the main functional entities in cells, the exclusive use of mRNA measurements comes at the risk of missing important biological information. Here we leverage recent technological advances in single-cell proteomics by Mass Spectrometry (scp-MS) to generate the first scp-MS dataset of an *in vivo* differentiation hierarchy encompassing over 2,500 human CD34+ hematopoietic stem and progenitor cells. Through integration with scRNA-seq, we identify proteins that are important for stem cell quiescence, which were not indicated by their mRNA transcripts, and demonstrate functional expression covariance during differentiation that is only detectable on protein level. Finally, we show that modeling translation dynamics can infer cell progression during differentiation and explain 45% more protein variation from mRNA than linear correlation. Our work serves as a framework for future single-cell multi-omics studies across biological systems.

## Introduction

Single-cell transcriptomics (scRNA-seq) of the human hematopoietic stem and progenitor cell (HSPC) compartment has changed our understanding of the hematopoietic system by revealing the gradual transition through cell transcriptome states during differentiation of HSPCs into functional blood cells^1–4^. Capturing enough of these transitional cell states from single-cell snapshots has enabled the indepth characterization of cell state heterogeneity and the recapitulation of differentiation trajectories^5,6^. Furthermore, modeling RNA velocity from mRNA splicing kinetics using scRNA-seq data has enabled the study of temporal dynamics during differentiation^7–9^, although its application to hematopoiesis has faced challenges due to time dependent kinetic rates^10,11^. However, albeit very successful, these methods have mostly been based on the transcriptome, while the actual workhorse for cellular function, the proteome, has not been widely accessible for single-cell analysis. The importance of proteome characterization has been shown in many multi-omics studies, encompassing mRNA and protein measurements on bulk population level, by revealing that transcript abundance can be a poor proxy for protein abundance^12–16^ and that the proteome can reveal more phenotypical information^17,18^. Furthermore, these bulk multi-omics datasets have enabled the modeling of translation kinetics during cell differentiation^16^, revealing the temporal dynamics between the two modalities^19–21^. On single-cell level, protein measurements have mostly been limited to targeted antibody-based cell surface epitope assays (e.g. CITE-seq), which were nonetheless instrumental to resolve cell states that were inaccessible with mRNA measurements^3,22,23^, and even enabled temporal modeling^24^. Thus, direct, untargeted and high-throughput measurements of proteins in single-cells have the potential to further improve our understanding of the hematopoietic system by providing more phenotypical information about observed cell states and by revealing temporal dynamics in the system.

Recent technological advances have enabled measuring protein expression in individual cells using single-cell proteomics by Mass Spectrometry (scp-MS)^25^. Specifically with a method termed SCoPE-MS^26^, where sample multiplexing via isobaric tags enables the combined measurement of 14-16 single-cells in a single mass spectrometry (MS) run. Importantly, the addition of a carrier channel provides more peptide ion copies to the MS to facilitate peptide identification. Based on this method, we have previously introduced our 384-well plate-based scp-MS workflow^27,28^, where single cells are isolated via fluorescence-activated cell sorting (FACS), which enables the incorporation of antibody-based surface markers to isolate or annotate populations of interest.

While scp-MS has reached a technological level that allows for the quantification of a significant part of the proteome, reports describing its application to *in vivo* differentiation hierarchies remain scarce. This poses the questions whether scp-MS can be scaled up for robust data generation of complex biological systems like the human HSPC compartment; and importantly, whether it captures phenotypical information that is complementary to mRNA-based approaches. This would make scp-MS well suited for single-cell multi-omics analyses, if additionally, integration of scRNA-seq and scp-MS data is demonstrated.

Here, we present a scp-MS dataset encompassing over 2,500 cells that recapitulates the HSPC hierarchy through relative protein expression changes. We subsequently computationally integrate scp-MS with scRNA-seq, generating a single-cell multi-omics dataset. We discover proteins involved in stem cell quiescence that are not apparent on mRNA level and highlight processes during lineage specification that are better described on protein level. Finally, we quantify temporal dynamics between the modalities and show that these can be modeled via a translation kinetics model that we termed scProtVelo.

## Results

### A single-cell proteomics dataset of FACS isolated human HSPCs

We isolated CD34+ bone marrow (BM) HSPCs from six healthy donors and stained the cells with a classical surface marker panel, enabling the isolation of defined subpopulations via FACS^2^ (**Figure 1A**). With the aim of measuring protein expression in single cells, we applied our scp-MS workflow^27,28^ on stained cells starting with the deposition of single cells into the wells of a 384-well plate, while recording the FACS parameters for later analysis (**Figure 1B**). We employed two different sorting strategies resulting in two kinds of multiplexed sets for scp-MS. First, we used the classical gating strategy to isolate specific subpopulations and distributed them evenly across the 14 single cells per multiplexed set (“Enriched Populations”). Secondly, we sorted any CD34+ cell resulting in the random sampling of the entire HSPC population (“Total HSPC”). For the MS measurement of each multiplexed set, we utilized our recently described real-time search-assisted acquisition (RETICLE)^28^. For downstream analysis, we employed our previously reported computational workflow SCeptre^27^, which expands on the functions of Scanpy^29^ for the processing of scp-MS data, including cell filtering, batch correction and data normalization (**Figure S1A-D**, see methods)

**Figure 1.**
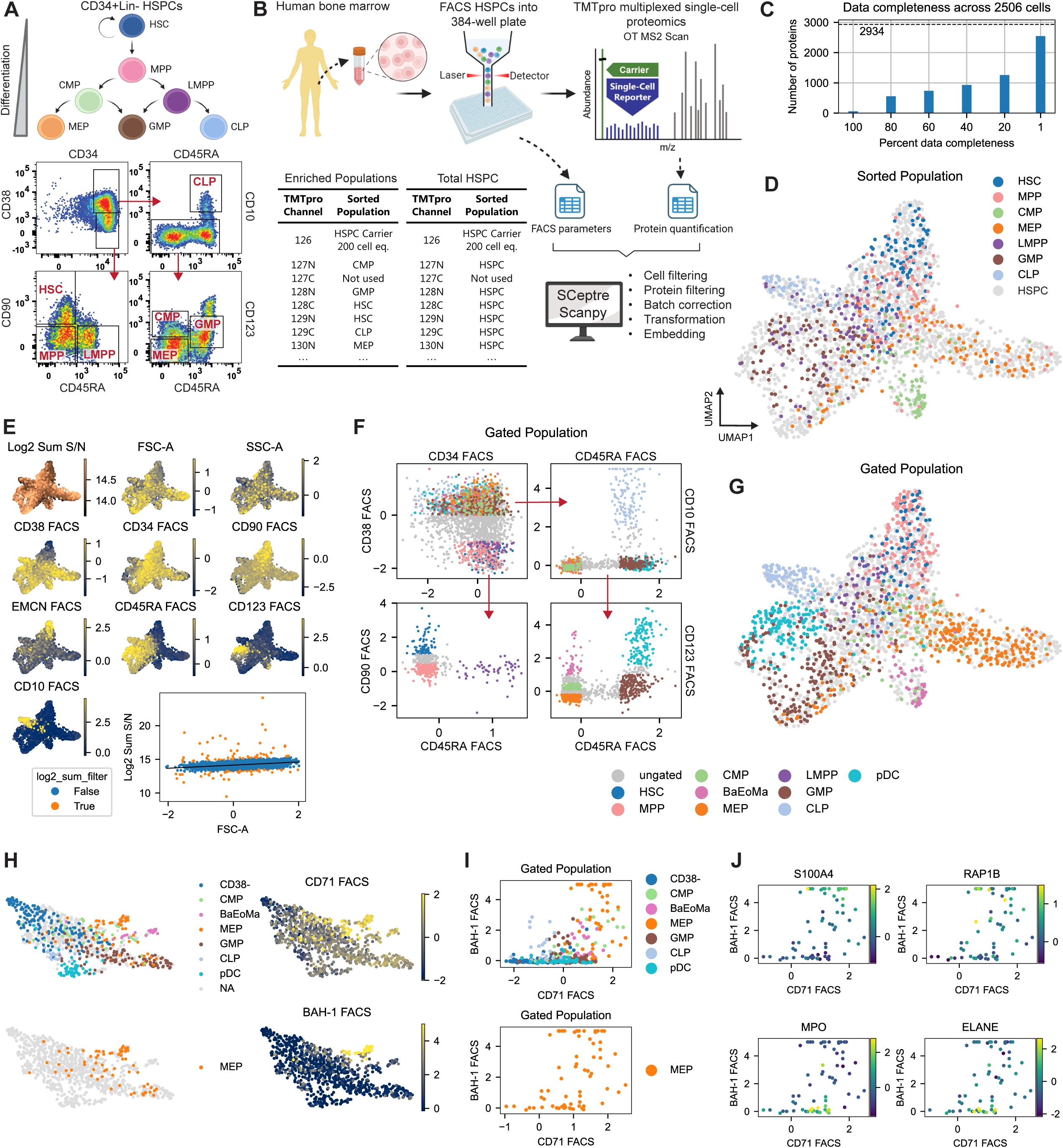
A single-cell proteomics dataset of FACS isolated human HSPCs. (A) The HSPC differentiation hierarchy (top) and sorting strategy (bottom). (B) Experimental workflow. Two different kinds of multiplexed sets were generated by sorting cells from specific gates (”Enriched Populations”) or any Lin-CD34+ cell (Total HSPC). (C) Data completeness of scp-MS dataset. (D) UMAP of cells from scp-MS data, annotated with their respective sorting gate. (E) Overlay of total MS signal and FACS parameters onto the UMAP. Scatterplot shows the relationship between FACS forward scatter area and scp-MS total signal. Orange outliers were removed during processing. (F) Computational gating strategy for retrospective annotation of cells from both enriched populations and total HSPC sorts. (G) UMAP annotated with gated populations from F. (H) UMAP of second scp-MS dataset with FACS markers enabling positive selection of MEPs. (I) Scatterplot showing the putative true BAH-1+CD71+ MEP subset. (J) Scatterplots from I with overlaid expression of MEP (top) and GMP (bottom) marker proteins. See also Figure S1.

SCeptre processing yielded a dataset with over 2,900 proteins quantified across 2,500 cells, albeit with variable data completeness (**Figure 1C**). From the UMAP^30^ embedding of the protein expression data, we did not observe any batch effects based on the TMTpro channel, individual donors or plates (**Figure S1B-D**), indicating that technical variations from the workflow and biological variations attributable to any specific donor sample were largely removed. We further validated this using principal component regression^31^ which revealed that only 4.6%, 0.71% and 0.02% of the variance could be explained by MS run, TMTpro label and donor, respectively. Overlaying the cell identities derived from the two different cell-sorting strategies onto the UMAP embedding revealed clustering and branching of the sorted (enriched) populations based on the HSPC differentiation hierarchy (**Figure 1D**). Hematopoietic stem cells (HSCs) and multipotent progenitors (MPPs) located at the top of the hierarchy and were followed by lymphoid-primed multipotent progenitors (LMPPs). LMPPs located upstream in the branches of granulocyte-macrophage progenitors (GMPs) and common lymphoid progenitors (CLPs), but not in the branch of megakaryocyte-erythroid progenitors (MEPs), consistent with their lineage output, which excludes the generation of MEPs^32,33^ (**Figure 1D**). Interestingly, we observed cells sorted as common myeloid progenitor (CMPs) to be clustered together with both myeloid branches but also forming a separate cluster that we later determined to be basophil, eosinophil, mast cell progenitors (BaEoMa) (see next section). This aligns well with findings that immunophenotypical CMPs are not a homogeneous population^34–36^ and with functional studies demonstrating BaEoMa potential within the immunophenotypic CMP population^37–39^. Taken together, these results showed that by processing the scp-MS data using the SCeptre pipeline, we successfully removed batch effects from technical covariates, without the need to define them. Since the SCeptre batch correction is based on alignments of medians and thus assumes balanced multiplexed sets, we expected good integration of the actively balanced multiplexed sets “Enriched Populations”. The observation that the batch correction also worked for the “Total HSPC” sets indicated that random samples of 14 HSPCs were sufficiently balanced.

Recording of the FACS parameters of each single cell during sorting enabled us to overlay that information onto the UMAP together with the total scp-MS protein signal (**Figure 1E**). This demonstrated the alignment of FACS markers and the scp-MS UMAP embedding. Cells differed in their scatter properties, indicative of differences in cell size and granularity. The differences in FSC-A correlated well with the total protein signal (**Figure 1E**), indicating a higher protein content for larger cells. Furthermore, our data confirms Endomucin (EMCN), a marker included in our antibody panel stain, to be specific for the most immature stem cells^40^. Therefore, our data suggests that Endomucin might serve as an alternative surface marker for enriching LT-HSCs, since commonly used markers CD90 and CD49f (data not shown) exhibited lower or no specificity in our dataset.

We subsequently used the recorded FACS parameters to perform retrospective computational cell gating (**Figure 1F**). We refined the gating strategy in the CD38+CD10-fraction (lower right panel) by moving the CMP gate down to CD45RA-CD123dim to incorporate a CD45RA-CD123+ gate for BaEoMa^39^. Furthermore, we moved the GMP gate to CD123-/dim to incorporate a CD45RA+CD123+ population of progenitors for dendritic cells (pDC)^2^. Overlaying these annotations onto the proteomics-derived embedding revealed well-clustered populations in line with the HSPC differentiation hierarchy (**Figure 1G**).

We observed that some MEPs clustered together with the GMP branch (**Figures 1D and 1G**). MEPs are purified based on the absence of specific markers; however, this population has been shown to be impure in functional assays^34,41^. To refine the MEP definition, we generated an additional scp-MS dataset with 922 cells, incorporating CD71 (transferrin receptor) and CD110/BAH-1 FACS markers to further specify the erythroid and megakaryocyte progenitors within the conventional MEP gate^34^ (**Figures 1H, S1E and S1F**). The combination of CD71 and BAH-1 clearly delineated MEPs, allowing the separation of those MEPs that clustered with GMPs (**Figures 1H and 1I**). Investigation of protein expression revealed proteins associated with MEP differentiation, such as S100A4 and RAP1B, to be highly expressed in the BAH-1 positive MEPs, and proteins associated with granulocytic differentiation, such as ELANE and MPO, to be highly expressed in some of the putative false MEPs lacking BAH-1 expression (**Figure 1J**). This observation corroborates the proximity of the BAH-1 negative cells in the MEP gate to the GMP branch.

Overall, our scp-MS data recapitulated the human HSPC hierarchy and highlighted certain limitations of low-dimensional FACS marker combinations in resolving the cellular heterogeneity.

### Unsupervised clustering reveals distinct cellular differentiation stages and functional protein covariance

Next, we explored the protein expression that drove the separation of HSPCs in this high-dimensional space that is not attainable with low-dimensional targeted surface marker assays. We used unsupervised cell clustering to group cells into 11 clusters (**Figure 2A**) that were named based on FACS information and protein expression (see this section). Next, we evaluated the quantitative accuracy of the scp-MS data by comparing fold-changes generated from cluster averages (i.e. pseudobulks) to bulk proteomics data derived from human HSPCs (HSC, MEP, CMP, GMP) measured via Data Independent Acquisition (DIA) LC-MS^42^. Thus, this dataset enabled both a comparison to bulk proteomics, and results from a different lab using a different MS method (**Figures 2B and 2C)**. For the comparison, we also included the computationally gated scp-MS populations, as they might better resemble the cell composition of the bulk-sorted populations. Pearson correlation values between protein log2 fold-changes of scp-MS-based populations and the four bulk populations revealed that scp-MS populations clustered together by similarity and had the highest correlation coefficients with their corresponding bulk population. The GMP and CMP bulk populations had slightly lower and much lower correlations respectively, likely due to minor differences between the bulk and single-cell FACS gating. Furthermore, we used the bulk data to benchmark cell expression normalization based on total signal versus median-ratio normalization and found the latter to perform slightly better (**Figure S2A**).

**Figure 2.**
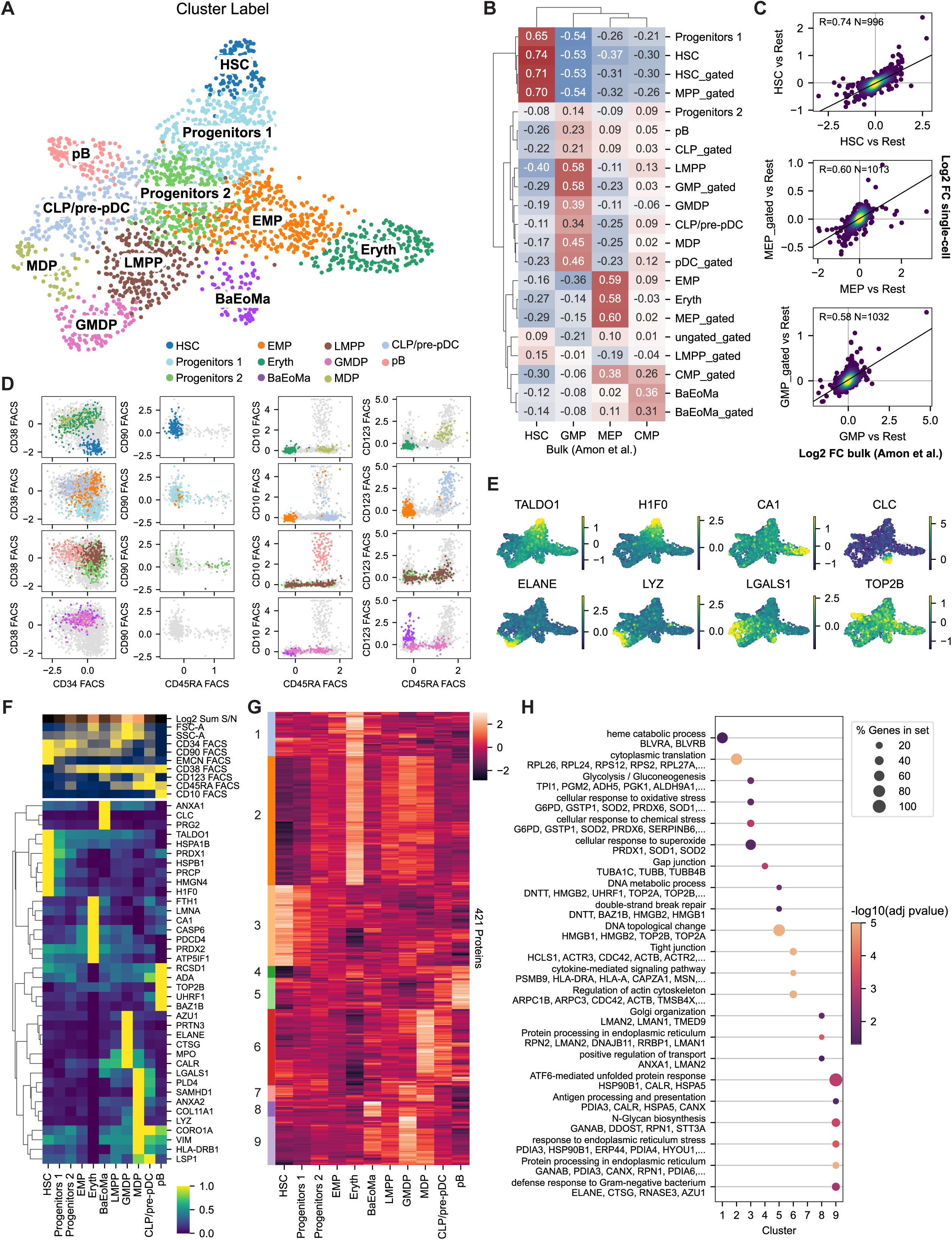
Unsupervised clustering reveals distinct cellular differentiation stages and functional protein covariance. (A) UMAP of cells annotated with leiden clusters. (B) Clustered heatmap showing the correlation of expression between scp-MS measurements and bulk proteomics data^42^, computed as Pearson correlation values of mean protein log2 fold-changes of a population vs all other populations. (C) Scatterplots of selected comparisons from B. (D) Cell clusters are visualized on FACS gating scheme from Fig.1F. Clusters were distributed over 4 panels for visibility. (E) UMAP annotated with the expression of selected proteins. (F) Clustered heatmap of selected cluster specific proteins. Mean aggregated and normalized between 0 and 1. Top shows aggregated FACS parameters. (G) Clustered heatmap of protein expression across leiden clusters. Mean aggregated. (H) Dotplot of top enriched terms per protein cluster. -Log10(adj. pvalue) was truncated to max. 5. See also Figure S2.

To evaluate the purity and integrity of the scp-MS clusters, we overlaid the cluster labels onto the computational gating scheme from Fig. 1F (**Figure 2D**). Most clusters were well restricted to their respective FACS gate. Only Progenitors 2 and, to some degree, Progenitors 1 were distributed over multiple FACS gates, indicating that these populations contain a mixture of differentiation stages. The erythroid-megakaryocyte progenitors (EMP) cluster should give rise to another branch containing megakaryocyte progenitors; however, we did not observe any putative branch, which might be due to their rarity in BM or their depletion through the inclusion of CD61 megakaryocyte marker in the lineage cocktail used for negative selection during FACS.

We next employed differential protein expression analysis to identify marker proteins for the 11 clusters (**Figures 2E and 2F)**. For example, HSCs showed high levels of H1.0 linker histone (H1F0), indicating a compact chromatin structure and quiescent cell state^43^. Erythroid differentiation was marked by the expression of Carbonic anhydrase I (CA1)^44^. Basophil-eosinophil-mast progenitors (BaEoMa) were marked by CD123 surface expression and protein expression of Proteoglycan 2 (PRG2) and Charcot-Leyden Crystal Galectin (CLC), components of eosinophil granules^45^. Granulocyte-monocyte-dendritic cell progenitors (GMDP) contained cells expressing the proteins AZU1, PRTN3, ELANE, CTSG and MPO that are found in azurophil granules, the first granules to be formed during neutrophil differentiation^46^. Progenitors for monocytes and common dendritic cells (MDP) branched off from the GMDP cluster, marked by the expression of LGALS1 and PLD4, which could also be observed in scRNA-seq data^47^ (**Figure S2B**). Furthermore, MDPs showed increased surface expression of CD123 and expression of proteins like Lysozyme (LYZ) and HLA-DRB1^3^. The cluster CLP/pre-pDC that likely contained a mixture of common lymphoid progenitors (CLP) and progenitors for plasmacytoid dendritic cells (pre-pDCs) shared markers with MDPs and progenitors for B-cells (pB). This is in line with the similarity of dendritic cell progenitors derived from myeloid or lymphoid lineages^48^, however, they could be differentiated from MDPs by the missing expression of e.g. LYZ^3^. Finally, the cluster that contained progenitors for B-cells (pB) was marked by surface expression of CD10 and protein expression of topoisomerase IIβ (TOP2B), which has been shown to be essential for B-cell development^49^. Thus, although HSPC differentiation is a continuous process, protein and FACS data are in good concordance on distinct expression changes during lineage commitment.

To investigate protein covariation across HSPC differentiation, we performed unsupervised clustering of the most variable protein expression profiles across clusters and performed gene ontology term enrichment analysis^50^ on each protein cluster (**Figures 2G and 2H**). Protein cluster 3 was highly expressed in HSCs and showed enrichment in glycolysis, likely reflecting their reliance on glycolysis instead of oxidative phosphorylation^51^. Furthermore, HSCs showed high levels of proteins related to the response to oxidative and chemical stress, crucial for protection from cellular damage. While cluster 3 expression gradually decreased during differentiation, cluster 2 gradually increased and was enriched in translation, reaching the highest levels in the erythroid lineage, but also high levels in GMDPs, indicative of the high translational activity needed to proliferate and to produce e.g. hemoglobin or granule proteins, respectively. Cluster 1 contained erythroid-specific proteins, e.g. related to heme processing. Cluster 5 was highly expressed in pB and enriched in terms related to the DNA rearrangements during B-cell development. Cluster 6 was highly expressed in MDPs and also elevated in CLP/pre-pDC and enriched in terms that are in line with macrophage and dendritic cell functions like motility, phagocytosis and antigen presentation. Cluster 9 was highly expressed in GMDPs and enriched in proteins contained in the azurophil granule and processes related to endoplasmic reticulum stress, e.g. due to the extensive production of granule proteins.

Overall, our observations capture a diverse range of biological processes, adding further validity to the biological relevance of our dataset and highlighting the potential to analyze protein covariation at scale using scp-MS data.

### Single-cell multi-omics analysis of transcriptomics and proteomics of the human HSPC compartment

With the aim to perform an integrated analysis of mRNA and protein expression in the HSPC compartment, we created a CITE-seq^22^ dataset of human HSPCs (**Figure 3A**). We integrated data from four individuals using totalVI^52^, resulting in a dataset of 9,086 cells. From a UMAP embedding of the mRNA expression data, we did not observe any batch effects based on specific individuals (**Figure S2C**), indicating that technical and biological variation attributable to any specific donor sample were largely removed. We annotated cells based on mRNA expression with a reference mapping from Azimuth^4^ and, similar to the scp-MS data, subsetted GMPs into GMDP and MDP (**Figure S2D**). The cell annotation in combination with the UMAP embedding, PAGA^53^ connectivity and surface marker expression indicated a good recapitulation of the human HSPC hierarchy (**Figures 3A, S2E and S2F**). Furthermore, while cell cycle phase prediction resulted in an expected phase distribution across differentiation stages (**Figure S2G**), it also revealed strong cell cycle dependent clustering of the UMAP embedding, which likely explains the observed, but unlikely, connectivity between Early Eryth and LMPPs in the same phase. We also observed a very wide range of total mRNA counts and detected genes per cell differentiation stage with HSCs having on average 3.45 times less counts and 2.1 times less detected genes compared to Late Eryth, in line with the low transcriptional activity in HSCs^54^ (**Figure S2H**). Since totalVI returns modeled and denoised mRNA expression values, similar to scVI^55^, we benchmarked the fold-change accuracy using the bulk mRNA data from Amon et al. and found that scVI-modeled and median-ratio normalized mRNA data showed higher correlation with the bulk data and similar centering around zero (**Figure S2I**). Nonetheless, we note that the scVI-modeled values should be used with caution as they represent only the mean of a modeled distribution with potentially high uncertainty.

**Figure 3.**
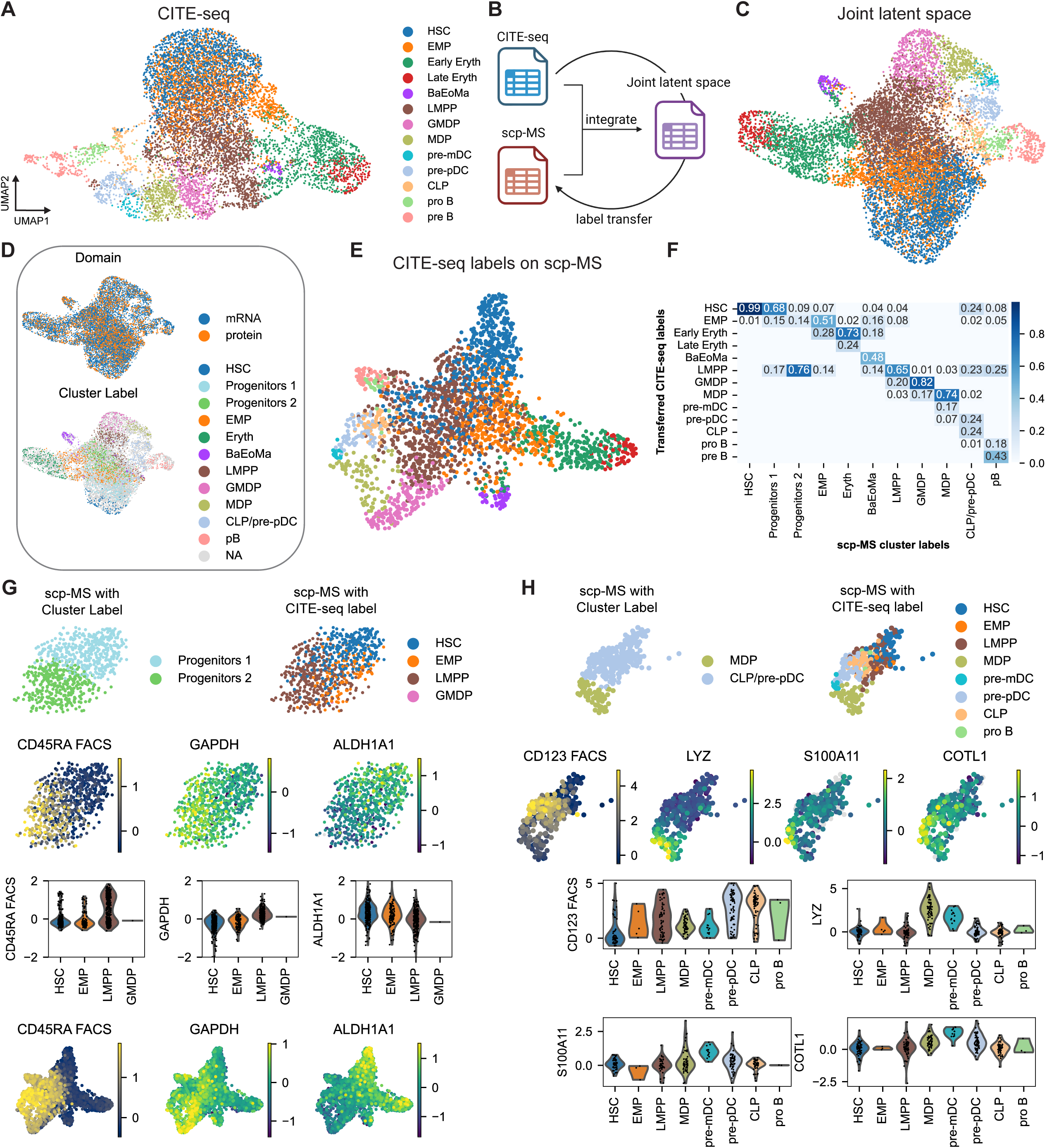
Single-cell multi-omics analysis of transcriptomics and proteomics of the human HSPC compartment. (A) UMAP embedding of annotated CITE-seq dataset. (B) Schematic of GLUE integration approach. (C) UMAP embedding of integrated dataset of CITE-seq and scp-MS, overlaid with CITE-seq cell annotation that was transferred to the “protein” cells. (D) Distribution of “mRNA” cells and “protein” cells (top). Overlaid cluster labels from scp-MS (bottom). (E) scp-MS UMAP embedding with transferred CITE-seq cell annotation. (F) Heatmap related to the label transfer from “mRNA” to “protein” cells. Rows are the cell annotation from CITE-seq, columns the cluster labels from scp-MS. The values represent the fraction of cells in the respective scp-MS cluster that was assigned the respective CITE-seq label. Thus, columns add up to one. (G) Validation of cell label transfer of scRNA-seq labels EMP and LMPP onto scp-MS labels Progenitors 1 and 2. (H) Validation of cell label transfer of scRNA-seq label pre-mDC onto scp-MS label MDP. See also Figures S2 and S3.

We integrated the scp-MS and CITE-seq datasets with GLUE^56^ (**Figure 3B**), a variational autoencoder (VAE)-based tool designed for unpaired single-cell multi-omics data integration. GLUE excels in this context by leveraging a user-provided guidance graph to link features across modalities, thereby preserving biologically relevant relationships without requiring paired or matched information. Specifically, we linked mRNA features from the CITE-seq data to their corresponding proteins in the scp-MS data. GLUE’s VAE framework is adept at handling noise inherent in single-cell data through probabilistic modeling, effectively distinguishing technical noise from biological signals. The resulting joint latent space successfully integrated “mRNA” cells from the CITE-seq dataset and “protein” cells from the scp-MS dataset. Both modalities overlapped almost perfectly with a silhouette score of 0.03. **Figure 3C** shows the UMAP embedding of the resulting joint latent space with “mRNA” cell labels transferred to the “protein” cells via nearest neighbor matching. Biological variation from the individual modalities was preserved in the integrated embedding as metrics on cell type separation^31^ were comparable before and after integration for both modalities (**Figure 3D and S2J**). Moreover, we found many protein-level marker to agree with mRNA expression (**Figure S3A**). Overlaying the new labels onto the protein UMAP revealed a more nuanced clustering (**Figure 3E**). Further, we compared the cell identity of “protein” cells based on the scp-MS clusters to the transferred labels, as we would expect these independently created annotations to agree in large. Indeed, as shown in **Figure 3F**, labels were not transferred across lineage branches. Furthermore, “protein” cells in scp-MS clusters Progenitors 1 & 2 were assigned EMP and LMPP labels, which suggested that the model learned to separate these lineages at an early stage based on protein expression. Indeed, for the cells in these clusters we found early differences in expression of surface marker CD45RA (FACS) and scp-MS-derived proteins like GAPDH, LCP1 and PSME1 that were expressed higher in LMPP and its descendants, while the proteins SOD2 and ALDH1A1 were expressed higher in EMPs (**Figures 3G and S3B**). Moreover, the Azimuth label pre-mDC was transferred onto “protein” cells contained in the MDP cluster. Indeed, these cells could be separated from MDPs and pre-pDCs by unique protein expression patterns of e.g. CD123 (FACS) and scp-MS-derived proteins like S100A11, COTL1 and especially Lysozyme (LYZ) (**Figure 3H**). Taken together, these analyses pointed towards a successful integration of mRNA and scp-MS data, and the resulting ability to transfer mRNA-derived labels onto “protein” cells facilitated the refinement of cell states in scp-MS data.

### Trajectory analysis on joint latent space recapitulates HSPC differentiation

We found that the joint latent space represented the biological latent cell states and we now asked whether we could identify differentiation trajectories and relevant driving factors, in particular when compared to using scRNA-seq alone as previously done. We therefore used the integrated latent space as input for a trajectory analysis using cellrank^5,6^. Here, we used the pseudotime kernel (**Figure 4A**) that infuses directionality into a k-NN graph to compute a transition matrix. The subsequently detected terminal states and the cell-specific fate probabilities indeed represented the expected differentiation trajectories in our HSPC dataset (**Figures 4A, S3C and S3D**). First, HSC-EMP-Early Eryth-Late Eryth for erythroid differentiation with early branching from HSCs into EMPs. BaEoMa displayed multiple origins from myeloid precursors, as observed previously^38,57^. Another trajectory was HSC-LMPP-GMDP with the end state enriched in precursors for granulocytic neutrophils. Alternatively, the pre-mDC trajectory shared the same start of HSC-LMPP-GMDP and then branched off to MDPs, which could branch off into monocytes (not part of HSPCs) or pre-mDCs. The lymphoid lineage for B-cell progenitors (pre B) appeared to have multiple origins from lymphoid progenitors, as also suggested by the Azimuth atlas (**Figure S3E**). Finally, the trajectory to pre-pDCs was HSC-LMPP-CLP-pre-pDC. We note that more trajectories might exist, as e.g. dendritic cell development has the potential for plastisticity^58^.

**Figure 4.**
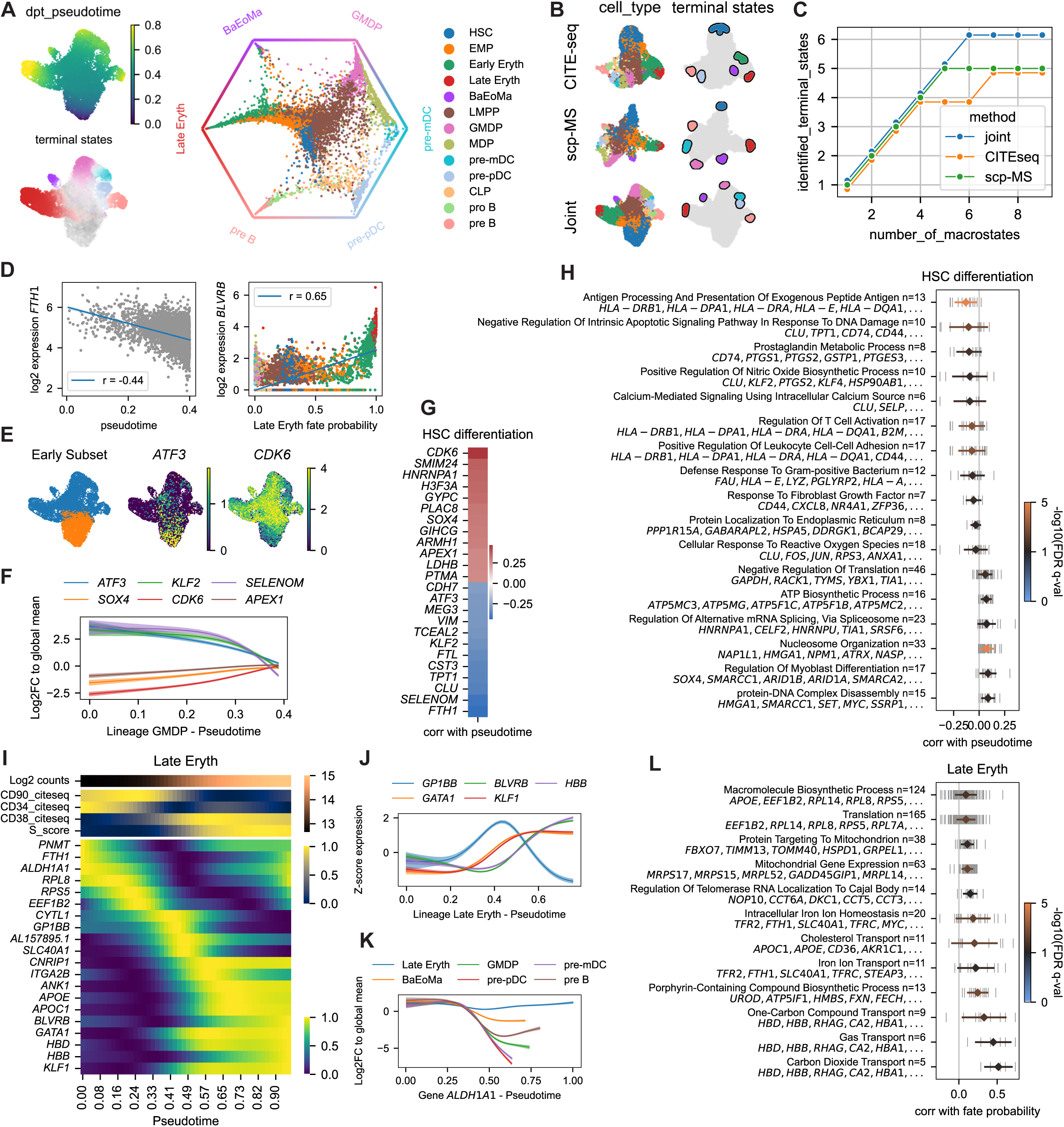
Trajectory analysis on joint latent space recapitulates HSPC differentiation. (A) UMAP with overlaid pseudotime and terminal states determined by cellrank (left). Cellrank circular projection showing cells ordered by their relative fate probability for each lineage (right). (B and C) Comparison of cellrank terminal state detection between CITE-seq, scp-MS and the joint dataset. B shows the top six detected terminal states. C shows the number of correct terminal states (y-axis) at different numbers of macrostates used for terminal state determination (x-axis). (D) Plots exemplify the calculation of correlation of gene expression with pseudotime (left) or fate probability (right). (E-G) mRNA expression analysis in an early subset of cells to investigate HSC differentiation. (E) Cells that were selected as early subset and mRNA expression of selected genes with high correlations. (F) Modeled gene expression profiles along pseudotime. The early part of the GMDP lineage was used for visualization. (G) Genes with the highest correlation or anticorrelation with pseudotime. (H) GSEA of HSC differentiation. X-axis shows correlation of expression with pseudotime. Vertical grey lines represent individual genes in the set. The horizontal bar represents the mean and standard deviation. (I) Heatmap showing mRNA expression profiles of selected genes in the Late Eryth trajectory. The S-phase cell cycle score was used to visualize the G1-S phase transition. (J) Modeled gene expression profiles of selected genes in Late Eryth lineage. (K) Modeled gene expression profiles of ALDH1A1 for the different lineages. (L) GSEA of Late Eryth trajectory. See also Figures S3 and S4.

We subsequently investigated the differences between the cellrank model based on the joint latent space and the CITE-seq and scp-MS datasets. We found that the terminal states identified from the joint latent space were not readily recovered from the CITE-seq or scp-MS datasets only (**Figure 4B**), even when more macrostates were considered (**Figure 4C**). Furthermore, we compared the fate probabilities across the three models, which revealed a better separation of the pre-pDC and pre-mDC trajectories in the joint latent space compared to CITE-seq (**Figure S3F**). Taken together, our results indicated that the cellrank model based on the joint latent space outperformed the models based on CITE-seq or scp-MS alone. These findings confirm our hypothesis that the two modalities provide complementary information about hematopoietic cell differentiation.

Subsequently, we used the results of the trajectory analysis to investigate mRNA and protein expression changes during differentiation. First, we analyzed HSC quiescence and differentiation by correlating mRNA or protein expression to pseudotime in an early subset of cells (**Figures 4D and 4E**). This yielded a correlation-based ranking of mRNA or protein, with positive correlation reflecting an expression increase during differentiation and negative correlation reflecting a decrease. Secondly, we analyzed the process of lineage specification by correlating mRNA or protein expression to fate probability for a particular lineage (**Figure 4D**). This yielded a correlation-based ranking of mRNA or protein, with positive correlations reflecting a coordinated increase of expression across single-cells during differentiation into that lineage compared to all other lineages. Importantly, the cell feature vectors pseudotime and fate probability were derived from the joint latent space and thus enabled the comparison of mRNA and protein expression by comparing the correlation vectors between modalities. We envisioned this analysis approach to be suitable for biological systems characterized by gradual transitions as in HSPCs, in contrast to a cluster-based analysis, where fold-changes between pseudobulk clusters are compared between mRNA and protein level. The correlation-based analysis does not require clustering and is computed on single-cell level; emphasizing features that are changing along the axis of interest (e.g. pseudotime or fate probability). And with the fate probabilities from cellrank, it makes full use of the soft lineage assignments per cell, emphasizing lineage specific changes.

To show the validity of this approach, we first analyzed only the mRNA expression using the “mRNA” cells. The analysis of HSC quiescence and differentiation revealed genes associated with HSCs like the transcription factors *ATF3*, which has been shown to prevent HSC exhaustion^59^, or *KLF2*, which is important for embryonic stem cell self-renewal^60^ (**Figure 4E-G**). In contrast, genes associated with stem cell differentiation were *CDK6*, a key regulator of exit from quiescence^61^, the transcription factor *SOX4*^62^ and the endonuclease *APEX1*^63^, which are both important for progenitor formation. Additionally, we modeled these gene expression profiles along the pseudotime to enable a visualization (see methods and **Figure 4F**). Subsequently, we used the correlation vector to perform a pre-ranked gene set enrichment analysis (GSEA)^50,64^ (**Figure 4H**). This revealed known processes for HSC maintenance such as antigen presentation via MHC II^65^, prostaglandin metabolism^66^, nitric oxide metabolism^67^ and reduction of reactive oxygen species Processes enriched during HSC differentiation were e.g. nucleosome organization and increased expression of the ATP synthase, required for the switch from glycolysis to oxidative phosphorylation^51^. Taken together, these results indicated that the pseudotime represented the main axis of early stem cell differentiation and supported the validity of our correlation-based analysis.

Continuing with the analysis of lineage specification, ordering those genes with the highest correlation for a lineage by their peak in pseudotime revealed a lineage-specific gene expression cascade (**Figures 4I, 4J and S4**). For example, the Late Erythroid trajectory was characterized by a transient expression of glycoprotein Ib platelet subunit beta (*GP1BB*), related to the potential branching into platelets forming megakaryocytes. Furthermore, we detected the expression of crucial erythroid differentiation factors *GATA1* and *KLF1*^68^ and subsequent expression of genes related to hemoglobin production (*BLVRB* & *HBB*). Additionally, this analysis revealed genes that separated a lineage by the loss of expression in other lineages, e.g. *ALDH1A1* expression was retained in the Late Erythroid trajectory (**Figure 4K**), as we have also observed on protein level (see **Figure 3G**). Again, we applied GSEA on the correlation vector, which revealed the enrichment of sets related to erythroid differentiation like iron transport and heme biosynthesis (**Figure 4L**). The remaining trajectories were visualized in **Figure S4**. For the BaEoMa lineage, we detected e.g. the transcriptional regulator *LMO4*^69^. The GMDP lineage was characterized by the early separation of LMPPs from EMPs marked by *SMIM24* and *C1QTNF4*^4^ and subsequent expression of granule genes (*AZU1*, *ELANE*). The pre-mDC and pre-pDC lineages were both marked via *LGALS1* and *SAMHD1,* while they could be separated by higher *IGSF6* expression in pre-mDC and higher *SPIB* expression in pre-pDC^4^. Finally, the pre B trajectory is marked by expression of *LAT2* and parts of the heterodimeric B-cell antigen receptor (*CD79A* and *CD79B*)^4^. Taken together, these results indicated that the cellrank model recapitulated correct differentiation trajectories of the HSPC hierarchy and supported the validity of our correlation-based analysis.

### Protein level information reveals additional insights into hematopoietic stem cell quiescence and differentiation

Subsequently we continued with the analysis of protein expression and directly compared the results to mRNA, by comparing correlation vectors between both modalities. For mRNA, we used the above calculated vectors based on counts, but also included correlation vectors based on the modeled and denoised scVI data. For protein, we repeated the analysis on the “protein” cells. The analysis of HSC quiescence and differentiation recapitulated the HSC protein markers we observed in **Figure 2F**, while proteins associated with HSC differentiation were mainly related to DNA replication. Interestingly, on mRNA level, many of these genes showed only low correlations or even disagreements in the direction of change in expression (**Figure 5A**). Comparison between the full correlation vectors on mRNA and protein level revealed an overall weak correlation below 0.25 between these vectors (**Figure 5B**). Moreover, this difference in the correlation vectors also resulted in the enrichment of gene sets that were not enriched on mRNA level (**Figure 5C**). For example, we observed an enrichment of proteins involved in carbohydrate metabolism (glycolysis) in HSCs, which was not enriched on mRNA level, in line with typically low correlations of key metabolic pathways^14^. Moreover, we found many proteins involved in the protection of HSCs from oxidative stress, including superoxide dismutases SOD1 and SOD2, Peroxiredoxin 1 (PRDX1), and TALDO1, a key enzyme of the pentose phosphate pathway that generates NADPH, which fuels the regeneration of reduced glutathione, a major cellular antioxidant^70^. Apart from SOD1, a clear decrease of their levels during early HSC differentiation could only be detected via scp-MS and not scRNA-seq (**Figure 5B**). We also found many proteins involved in chromatin structure among HSC-correlating factors, including the previously mentioned H1 linker histone H1F0, the H1-like protein HP1BP3, which is required for HSC self-renewal^71^, the histone macroH2A1 (H2AFY), associated with HSC homeostasis^72^, and the chromatin regulator HMGA1, important for HSC regenerative capacity^73^. Again, the decrease of these chromatin regulators during early HSC differentiation was better described via scp-MS. Finally, we observed strong differences between mRNA and protein levels of nuclear lamin B2 (LMNB2). It has been shown that Lamin-A:B stoichiometry regulates nuclear stiffness^74^, thus, scp-MS could potentially better inform about the nuclear morphology in single cells than scRNA-seq.

**Figure 5.**
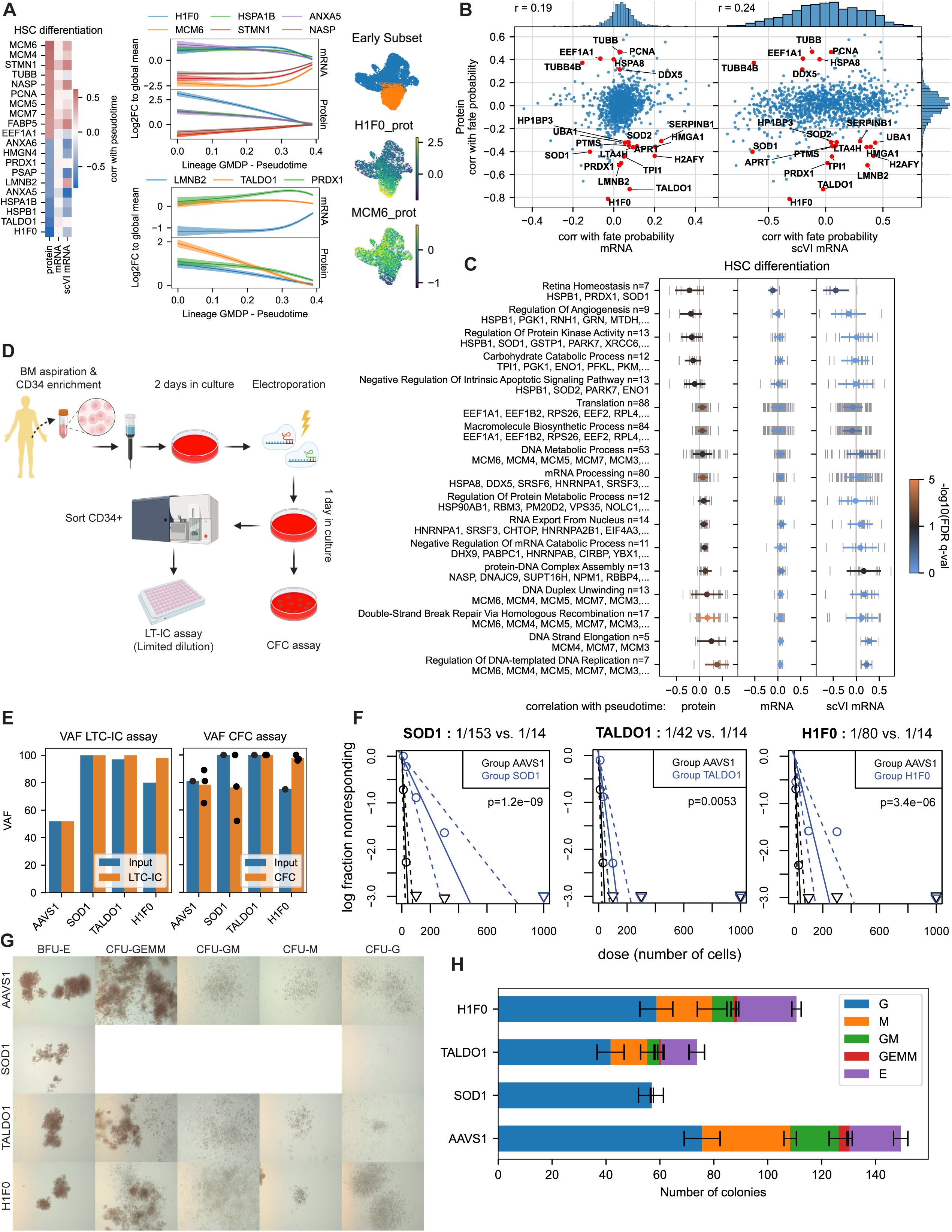
Hematopoietic stem cell quiescence and differentiation is described differently on protein level. (A) Protein expression analysis in an early subset of cells to investigate HSC differentiation. Left: Proteins with the highest correlation and anticorrelation with pseudotime including mRNA level information. Middle top: Modeled protein and mRNA expression of selected genes. GMDP lineage was used as an example. Middle bottom: Modeled protein and mRNA expression of selected genes that are not HSC-specific on mRNA level. Right: Cells that were selected as early subset and expression of selected proteins. (B) Scatterplot comparing the correlation vectors between protein and mRNA (count-based and scVI modeled). Selected genes were annotated. (C) GSEA analysis of correlation vectors of HSC differentiation. Vertical grey lines represent individual genes in the set. The horizontal bar represents the mean and standard deviation. (D) Experimental workflow functional assessment of selected proteins. (E) Variant allele frequency (VAF) of CRISPR knockout. (F) Quantification of LTC-IC limiting dilution assay. (G) Representative images of colonies of CFC assay. BFU=Burst Forming Unit, CFU=Colony Forming Unit, E=Erythroid, GEMM=Granulocyte/Erythroid/Megakaryocyte/Macrophage, GM=Granulocyte/Macrophage, M=Macrophage, G=Granulocyte (H) Quantification of CFCs. See also Figures S5 and S6.

Finally, investigation of the proteins and their gene sets related to HSC differentiation revealed processes that were better described via scp-MS. For example, the increase of proteins involved in mRNA processing, RNA export from the nucleus, translation and DNA replication (**Figure 5C**).

To validate the functional importance of selected proteins associated with the HSC state that have previously not been interrogated in human, we performed CRISPR/Cas9-mediated knockout of SOD1, TALDO1 and H1F0 in human CD34+ BM cells (**Figure 5D**). Subsequently, we conducted a long-term culture-initiating cell assay (LTC-IC) as a surrogate for assessing HSC activity upon knockout (**Figure 5D-F**). LTC-IC limiting dilution analysis^75^ revealed a reduction in LTC-IC frequency in SOD1 knockout cells (1 in 153) compared to the AAVS1 control (1 in 14), indicating a decreased ability to sustain long-term hematopoiesis upon knockout (**Figure 5F**). Furthermore, SOD1 knockout resulted in fewer colony-forming units (CFUs), with the remaining very small colonies predominantly displaying a granulocytic outcome (CFU-G) (**Figure 5G and 5H**). These findings not only demonstrate the importance of SOD1 for immature HSPCs, but also highlights species differences between mice and man, as *Sod1 null* mice only display minor hematopoietic phenotypes^76^. The knockout of TALDO1 and H1F0 resulted in a reduced stem cell frequency of 1:42 and 1:80, respectively (**Figure 5F**). While there was an overall decrease in the number of CFUs in TALDO1 and H1F0 knockout, the proportion of multilineage output remained unchanged (**Figure 5G and 5H**), suggesting a specific impact on the most immature cells.

In summary, the protein expression trends during early HSC differentiation revealed additional information compared to our scRNA-seq measurements and identified novel proteins and related pathways important for HSC function.

### Proteomics-based trajectory analysis reveals protein level-specific functional covariation

Besides maintenance of the HSC state, we asked which protein-level programs are specific to particular lineages. For this, we analyzed lineage specification on protein level in the integrated dataset and compared it directly to the mRNA level (**Figures 6A-G, S5 and S6**). This revealed many of the protein markers from **Figure 2F** but also new proteins. For example, the Late Erythroid trajectory was characterized by the retained expression of ALDH1A1 (**Figures 6A and 6C**), as described above. Furthermore, we detected the transient expression of markers for megakaryocyte progenitors like RAP1B^4^, and subsequent expression of erythroid related proteins like S100A4, PRDX2 and CASP6 (**Figures 6A and 6C**). Comparison between the full correlation vectors on mRNA and protein level for this lineage (**Figures 6D and S5C**) revealed a much higher agreement of up to 0.7 compared to the HSC differentiation (r=0.19-0.24). Nonetheless, differences between mRNA and protein correlation vectors propagated to differences in the GSEA, including e.g. processes involving ribosomal proteins (**Figure S6**). In the GMDP lineage (**Figures 6B, 6C and 6E**), we found gene sets related to protein glycosylation and endocytosis to be enriched only on protein level (**Figure S6**). The latter set contained Beta-2 microglobulin (B2M), which, in our dataset, was not well correlated between mRNA and protein, since e.g. scp-MS measured an increase of B2M in the GMDP lineage, while the mRNA measurement indicated lower relative expression levels (**Figures 6E and 6F**). Nonetheless, the B2M protein expression profile corresponded well to its complex partners, HLA-A and HLA-B in the major histocompatibility complex (MHC) class I (**Figure 6F and 6G**).

**Figure 6.**
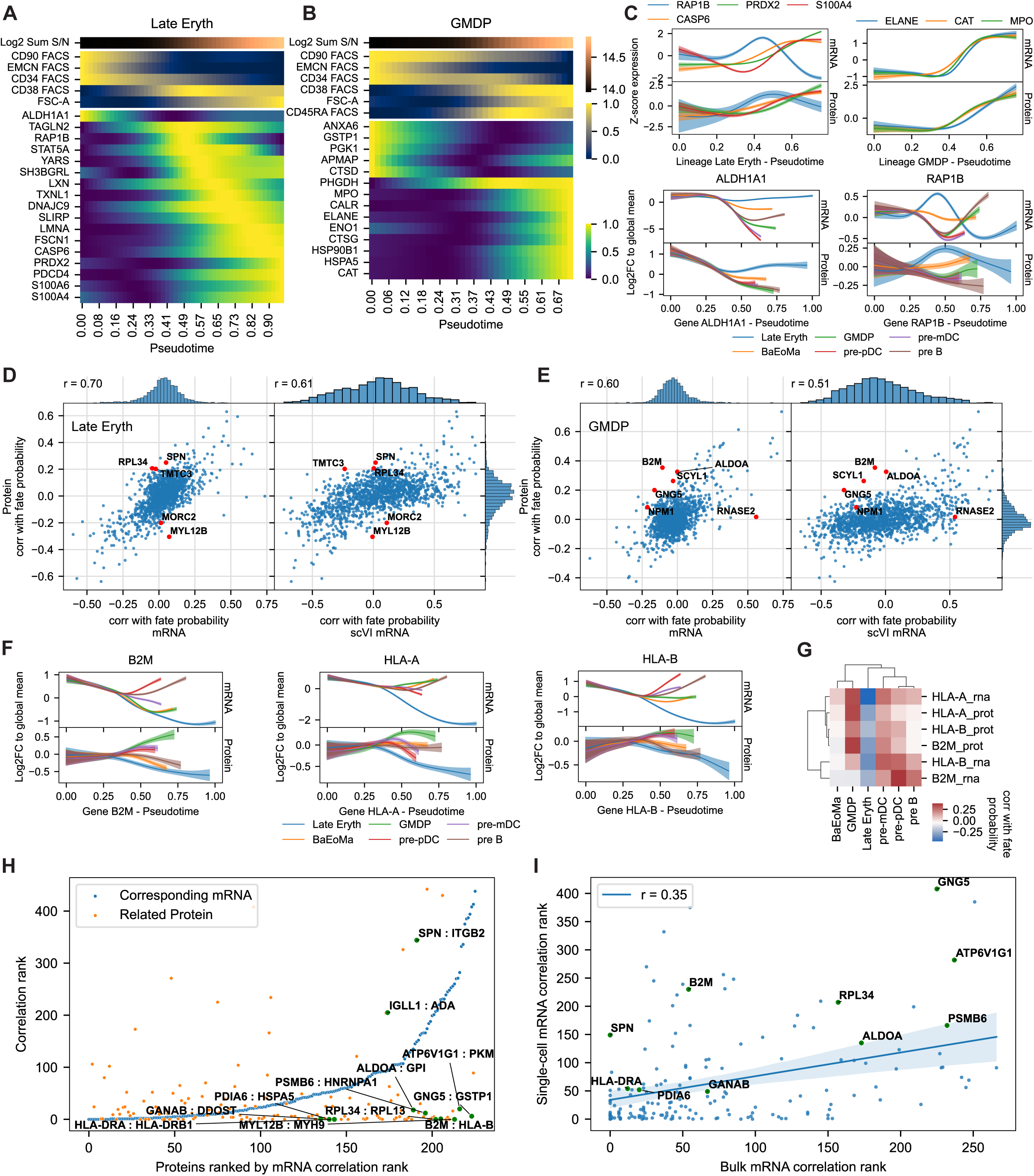
Protein level trajectory analysis reveals protein level-specific functional covariation. (A and B) Heatmap showing expression profiles of selected proteins in the Late Eryth and GMDP trajectories. (C) Modeled expression profiles on protein and mRNA level of selected genes. (D and E) Scatterplot comparing the correlation vectors between mRNA and protein level in the Late Eryth and GMDP trajectories. (F) Modeled expression profiles of MHC I complex members B2M, HLA-A and HLA-B. (G) Clustered heatmap of expression profiles of MHC complex members on both mRNA and protein level. (H) Scatterplot of proteins (x-axis) and the correlation ranks of both mRNA and protein molecules (y-axis). Proteins (x-axis) were sorted by the correlation rank of the corresponding mRNA (blue dots). Orange dots depict the related protein (same KEGG pathway) with the lowest correlation rank. (I) Comparison of mRNA correlation rank between bulk and scp-MS data.

Based on these results, we hypothesized that the proteins with low concordance with their respective mRNA template still covaried with functionally related proteins. To investigate this, we performed a joint mRNA-protein covariation analysis. Since the trajectory analysis yielded six different lineages, for each mRNA and protein molecule, the six correlations with lineage fate probabilities resulted in a vector of six values, representing the relative expression level across lineages. We focused on the 226 most variable proteins and their respective mRNA transcript and calculated a correlation matrix between those 452 molecules. Then, for each protein, we ranked all molecules by correlation and calculated the rank of the corresponding mRNA. We found that for a large fraction of proteins, the respective mRNA molecule ranked very high, indicating that it was among the best predictors for the protein expression profile across lineages (**Figure 6H**, blue dots). Subsequently, for each protein, we calculated the rank of the highest-ranking protein that was part of the same KEGG pathway. This revealed that many protein expression profiles could be well explained by a functionally related protein (**Figure 6H**, orange dots). Furthermore, looking at the proteins with low mRNA concordance, i.e. high mRNA ranks (right side of the plot), we indeed found that many protein expression profiles could be better explained by functionally related proteins than their respective mRNA (**Figure 6H**). Apart from the B2M-HLA-B pair, more examples were ALDOA and GPI, both part of glycolysis, MYL12B and MYH9, both part of the myosin complex, HLA-DRA and HLA-DRB1, both part of the MHC class 2 complex, and lastly, protein pairs involved in protein processing in the endoplasmic reticulum GANAB : DDOST and PDIA6 : HSPA5 (**Figure 6H**). Finally, to exclude that the observed low concordances between mRNA and protein was not purely due to dataset specific measurement errors, we compared our results to an external bulk dataset with mRNA and protein measurement of 59 breast cancer cell lines^77–79^. Calculating the mRNA correlation rank for each protein in this dataset revealed a positive correlation of 0.35 with our ranks (**Figure 6I**). These results supported a general trend for some genes to exhibit low correlations between mRNA and protein^13,14^, however also indicated that there might be effects that are specific to the biological system or measurement technique.

### scProtVelo models translation dynamics in erythroid differentiation

Finally, we asked whether we could connect protein kinetics to mRNA expression via a translation model, implying a delay in protein expression changes that could explain low mRNA-protein correlation values (**Figure 7A**). For this, we developed scProtVelo (single-cell protein velocity) to infer temporal gene expression dynamics from paired single-cell mRNA and protein expression data. scProtVelo jointly models both expression levels over time via their translation dynamics described by gene-specific transcription, translation, mRNA degradation and protein degradation rates (**Figure 7B**). We built the scProtVelo implementation onto the veloVI framework^9^ originally set up to model splicing dynamics for RNA velocity estimation. While scProtVelos translation modeling is conceptually similar to RNA velocity, there are also some key differences. While unspliced and spliced mRNA expression levels follow conservation of mass, mRNA molecules are not converted into protein molecules. Also, while splicing dynamics seem to have an initial state where both unspliced and spliced expression of a gene are zero, we observed the need for more flexible modeling of initial states. And lastly, we observed more accurate modeling, when we restricted the dynamic state (activation or repression) to be fixed per gene instead of enabling a switch in transcription rate within the modeled data. scProtVelo uses variational inference to fit a probabilistic formulation of the described translation dynamics to the data. The resulting variational autoencoder architecture takes a cell representation as input and infers a time annotation together with gene-specific, learnable transcription, translation as well as RNA and protein degradation rates, thus reconstructing the original mRNA and protein expression levels. Per gene we fit both dynamic states (activation and repression) simultaneously to the data as well as a Dirichlet distribution that represents the state assignment per gene (**Figure 7C, Methods**). We created paired measurements that can be modeled by scProtVelo by interpolating expression values of the missing modality of a cell from neighboring cells using the joint GLUE neighborhood graph and directly use the latent cell representations as input to the scProtVelo encoder.

**Figure 7.**
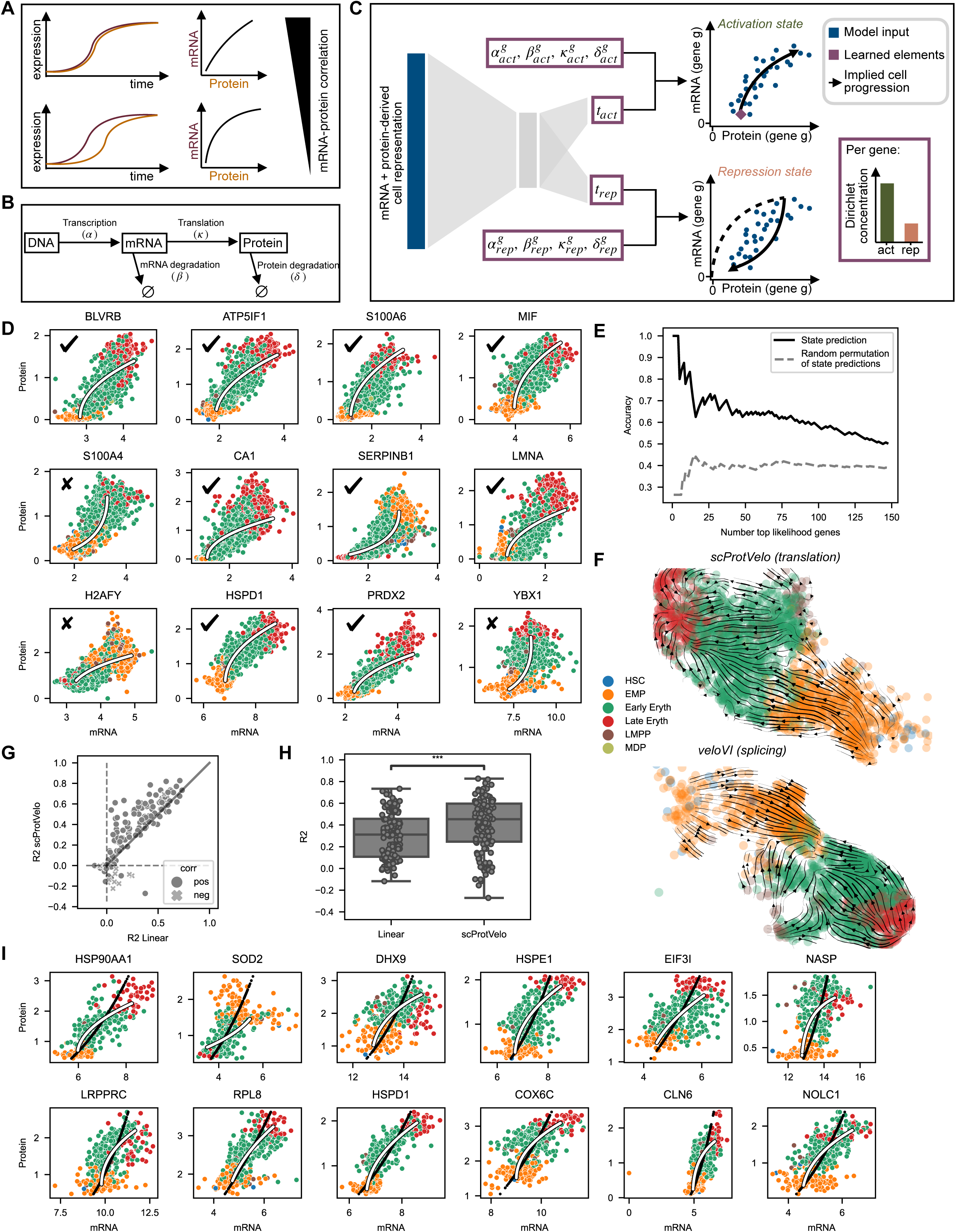
scProtVelo models translation dynamics in erythroid differentiation. (A) Expression time delay decreasing correlation. (B) Translation dynamics governing mRNA and protein expression. (C) Schematic of the scProtVelo architecture. (D) Phase portraits of the genes with highest model likelihoods. Ticks indicate genes where model fits agree with true cell progression. (E) Performance of gene state predictions compared to uninformed predictions on genes that significantly change along the selected trajectory. (F) Translation-based (top) and splicing-based (bottom) velocities derived from the top 100 likelihood genes projected onto a UMAP embedding. (G) Gene-wise comparison of protein expression variance explained by scProtVelo and a linear model, differentiated by the sign of the linear regression slope. (H) Global comparison of the protein expression variance explained by scProtVelo and a linear model. (I) Phase portraits showing the difference between fits from scProtVelo (white) and a linear model fitted in log space (black). See also Figure S7.

Since we start with a set of cells without corresponding time measurements or information about the direction of change of a gene over time, the only element that can inform the model about the underlying dynamics is the time delay that we expect between mRNA and protein expression changes. So, we first set out to test whether scProtVelo can pick up enough signal to infer correct cell transitions and gene states within our differentiation data. To have some ground truth, we subsetted our dataset to the erythroid differentiation trajectory and compared inferred dynamic gene states to differential expression results between early and late cells. We indeed found that time delay in expression was detectable, as model predictions were significantly better than random permutations of the model predictions (**Figures 7D and 7E**). Sorted by unsupervised model likelihood, state predictions reached an accuracy of 73% within the top 26 genes (**Figure 7E**). Further, aggregating the inferred protein velocity information of the top 100 genes recapitulated the correct cell progression from EMPs to Early and then Late Erythroids (**Figure 7F top**). In contrast, applying the standard RNA velocity workflow to the scRNA-seq cells resulted in the previously reported erroneous backflow in velocity vectors from

Late to Early Erythroid progenitors^10,11^ (**Figure 7F bottom**). Judging proposed cell to cell transitions by whether they coincide with an increase in pseudotime consolidates higher accuracy in trajectory inference based on translation modeling than RNA velocity (**Figure S7**).

Subsequently, we went back to the original question of how much mRNA-protein correlation can be increased by accounting for delay in expression. To quantify this, we considered 147 genes with significant variation during erythroid differentiation and incorporated pseudotime annotations as a prior on the inferred time into scProtVelo to ensure correct model fitting. Comparing the protein modeling abilities of scProtVelo with protein prediction results from a linear model, we found that scProtVelo resulted in more accurate modeling for almost all genes (except anti-correlated mRNA-protein-pairs) (**Figure 7G**). Overall, accounting for translation dynamics when modeling protein expression from mRNA levels leads to a 45% increase in explained protein variance as compared to the simple assumption of a linear relationship (median R2 values of 31% and 45% for the linear model and scProtVelo, respectively, **Figure 7H**). Genes with the strongest difference in explained variance between the modeling approaches are shown in **Figure 7I**.

Taken together, our results exemplify the successful modeling of temporal gene expression dynamics from integrated scRNA-seq and scp-MS data, its application for protein velocity-based trajectory inference and its contribution to illuminate the often low mRNA-protein correlations. scProtVelo is available as a python package via github.com/theislab/scProtVelo.

## Discussion

In this work, we presented the first scp-MS dataset of the primary human HSPC compartment, encompassing over 2,500 cells that recapitulated the HSPC hierarchy through relative protein expression changes. The combination with FACS measurements allowed us to validate the scp-MS-defined cell states and revealed heterogeneity in surface marker-defined populations. Furthermore, integration of scp-MS with CITE-seq data resulted in a joint latent space that enabled e.g. label transfer between modalities and a trajectory analysis that, in our hands, showed better performance compared to the two single modalities. Vectors derived from the joint latent space as e.g. pseudotime or fate probabilities enabled us to investigate differences between the modalities without the need for paired data. Here we found that the proteome level better described certain processes during HSC quiescence and differentiation, and we identified proteins which we experimentally validated to be important for the functional behavior of the most immature HSPCs.

In contrast to the HSC state, we found that the concordance between mRNA and protein is much higher during lineage specification, which possibly could be partly explained by the higher effect sizes resulting in increased signal-to-noise measurements. Nonetheless, we were able to show that many of the proteins were not well explained by their mRNA profiles. Further, many of these proteins co-varied with functionally related proteins, exemplifying the relevance of the protein expression profile. Future experiments could elucidate which post-transcriptional regulation mechanisms are at play. For example, the abundance of Beta-2 microglobulin (B2M) could be regulated on the protein level by protection from degradation via stabilization in the MHC I complex^80^.

Finally, we investigated the temporal dynamics between the modalities by modeling translation dynamics during erythroid differentiation. We found that this model could better explain the relationship between modalities than a linear regression, highlighting its relevance for shedding light on the often low correlations between the modalities. Further, we successfully used the learned dynamics to infer directionality of the erythroid differentiation process, which has been challenging to achieve using splicing kinetics^10,11^.

In the present work, we have generated and analyzed the first scp-MS dataset of an *in vivo* differentiation hierarchy and integrated it with mRNA-based readout. While technological advances in scp-MS will continue to improve data depth and accuracy in the future, our work serves as a framework for more single-cell mRNA/protein-based multi-omics studies. Such studies have the potential to improve our understanding of cell states and translation dynamics across many biological systems including normal development, stem cell systems, and disease states such as cancer.

## Code and data availability

Code and data will be made available with the publication of this manuscript. The scProtVelo package and analysis is available on github via github.com/theislab/scProtVelo.

## Acknowledgements

We thank Philipp Weiler for the setup of the first version of scProtVelo. B.F. is the recipient of a Copenhagen Bioscience Ph.D. stipend by the Novo Nordisk Foundation, NNF19SA0035442). S. R. is supported by the Helmholtz Association under the joint research school Munich School for Data Science. Work in the Porse lab was supported by grants from the Svend Andersen Foundation, the Candys Foundation, the Danish Cancer Society, the Eva and Henry Frænkel Memorial Foundation, the Independent Research Fund Denmark and through a center grant from the Novo Nordisk Foundation (Novo Nordisk Foundation Center for Stem Cell Biology, DanStem; Grant Number NNF17CC0027852). This work has been performed in the context of the Danish Research Center for Precision Medicine in Blood Cancers funded by the Danish Cancer Society (R223-A13071) and Greater Copenhagen Health Science Partners. Work in the Schoof lab was supported by grants from the Independent Research Fund Denmark (2067-00053B), Danish Cancer Society (R324-A17978), Novo Nordisk Foundation (NNF21OC0071016), and the Lundbeck Foundation (R413-2022-869).

## Author contributions

N.Ü. performed scp-MS sample collection and preparation. B.F. performed scp-MS measurements.

D.B. performed CITE-seq sample preparation and measurement. A.W. performed CITE-seq raw data analysis. M.B.S. performed the CRISPR knockout experiments. S.R. and F.T. created scProtVelo modeling. H.H. contributed to data interpretation. B.F. and S.R. analyzed and interpreted the data.

K.G. and K.T.M. aspirated human bone marrow samples, provided methodology and contributed to data interpretation. F.T., E.S., B.T.P. Supervision. E.S, B.T.P. conceived the study. B.F., N.Ü., S.R. Writing - Original Draft. F.T., E.S., B.T.P. Writing - Review & Editing. All authors approved the final version of the paper.

## Supplemental information

Table S1. FACS antibodies used.

Table S2. CITE-seq antibodies used.

## Methods

### Human bone marrow processing

Bone marrow samples were collected from healthy donors following the standard protocol of the Department of Hematology, Rigshospitalet, with prior informed and written consent according to the Helsinki declaration under a protocol approved by the Danish National Ethics committee (1705391). Briefly, 15 mL of bone marrow was aspirated into a 20 mL syringe containing 5 mL of ice-cold sterile-filtered anticoagulant ACD, and immediately mixed to prevent clumping. The collected BM was then diluted 1:2 with PBS + 2mM EDTA + 0.5% BSA (PBS-EDTA) and filtered through a 70µM filter. Subsequently, 30 mL of the diluted BM was gently layered over 15 mL of Lymphoprep (ProteoGenix, Cat. 1114544) and centrifuged at 400g for 30 minutes at 20°C. The resulting supernatant, containing the interphase with the low-density mononuclear cells (LDMNCs), was transferred to a new tube. The LDMNCs were washed at a 1:1 ratio with PBS-EDTA, and the pellet was resuspended in 10 mL of PBS-EDTA. Cell counts were determined using a NucleoCounter, and the cells were frozen in FBS containing 10% DMSO overnight at -80°C before being transferred to liquid nitrogen storage. For thawing, cells were gently warmed in a water bath at 37°C for 2–5 minutes and then transferred to a 50 mL Falcon tube. 20 mL pre-warmed thawing medium (RPMI + 10% FCS + 100µg/mL DNAse (Roche, Cat. 11284932001) + 5mM MgCl2) was added dropwise to the cell suspension, followed by centrifugation at 300g for 10 minutes at room temperature. The supernatant was removed, and the cell pellet was resuspended in 10 mL of pre-warmed thawing medium and incubated for 15 minutes at 37°C. After the incubation, cells were counted and centrifuged.

### FACS staining and cell sorting

All FACS experiments included single-color controls using CompBeads (BD Biosciences, Cat. 552844) and Fluorescence Minus One (FMO) controls. Antibodies are shown in **Table S1**.

#### scp-MS samples

Thawed BM mononuclear cells were CD34+ enriched using CD34 MicroBeads (Miltenyi Biotec, Cat. 130-046-702) according to the manufacturer’s instructions. CD34+ HSPCs were subsequently incubated with a cocktail of biotinylated lineage antibodies and the purified, unconjugated Endomucin antibody for 30 min on ice. After incubation, cells were washed with PBS-EDTA and centrifuged at 100g for 10 min. Subsequently, cells were incubated for 30 min with Streptavidin-PECy5 and Goat anti-Rabbit IgG (PE), followed by another wash with PBS-EDTA at 100g for 10 min. Finally, the cells were incubated with the panel of monoclonal antibodies for 30 min. Similar as previously described^27,28^, after three washes with PBS to eliminate any remaining serum, cells were sorted on a BD FACSAria III instrument at a maximum speed of 500 events per second into Eppendorf twin.tec 384 LoBind plates containing 1 µL 20% TFE, 10mM TCEP, 40mM CAA, 50mM TEAB lysis buffer. Live cells were sorted based on the membrane-impermeable viability dye 7-AAD. The 100µm nozzle was used for all sorts into 384-well plates, and "Single Cell" and "4-way Purity" precision modes were selected for single cell and 500 cells/well carrier sorts, respectively. Immediately after sorting, plates were briefly spun, snap-frozen on dry ice, and then boiled at 95°C in a thermocycler (Applied Biosystems Veriti 384-well) for 5 min. The plates were then briefly spun again and stored at -80°C until further sample preparation.

#### CITE-seq samples

BM cells were incubated with the cocktail of biotinylated lineage antibodies and the panel of monoclonal antibodies for 30 min on ice. Live cells were sorted based on the membrane-impermeable viability dye 7-AAD. Cells were FACS-sorted as Lin–CD34+ into low bind Eppendorf tubes using a BD FACSAria III instrument.

### scp-MS sample preparation

Plates were thawed on ice and sonicated for 2 min in a water bath sonicator. Protein lysates from the single cells were digested with 2 ng of Trypsin (Promega, Cat. V5280), dissolved in 1 μl of 100 mM TEAB pH 8.5 containing Benzonase (Sigma, Cat. E1014) diluted to a ratio of 1:5000 (vol/vol) to digest any DNA that would interfere with downstream processing. For the carrier plate, containing 500 cells per well, the trypsin amount was increased to 10 ng in each well. All dispensing steps in this procedure were carried out using the Dispendix I-DOT One instrument. After dispensing the digestion buffer, the plates were spun down, vortexed and incubated at 37°C overnight to complete the digestion.

On the next day, peptides were labeled with TMTPro 16plex reagents (ThermoFisher Scientific, Cat. A44520). 6 mM of each label (except 127C) in 1 µL acetonitrile (ACN) was added to the single cell wells, while the 500-cell carrier wells were labeled with 13 mM of TMTPro-126 reagent in each well. The plates were then incubated at room temperature for 1 hour. The labeling reaction was quenched by adding 1 μl of 1.25% Hydroxylamine (ThermoFisher Scientific, Cat. 90115) in 100 mM TEAB to single cell wells and 2.5% to carrier wells, followed by 15 min incubation at room temperature. The carrier plate, containing peptides labeled with TMTPro-126, was pooled into one sample and desalted using a SOLAµ Solid Phase Extraction (SPE) column (ThermoFisher Scientific, Cat. 60209-001) following the standard cleanup protocol. Eluted and desalted peptides were concentrated to dryness in an Eppendorf Speedvac, after which they were resuspended in A* (2% ACN, 0.1% TFA). Subsequently, peptides from 14 single cells and 200-cell equivalent of the Lin-CD34+ carrier were pooled and desalted using a SOLAµ SPE 96-well plate, described in the following. Briefly, the SOLAµ plate was prepared by activating with 200 µL methanol, conditioning with 200 µL 80% ACN, 0.1% FA, and equilibrating with 200 µL 0.1% FA. Between each step, the plate was centrifuged at 700g for 1 min. Next, 60 µL of 0.1% FA and 200-cell equivalent of the carrier (in 10 µL volume) were dispensed into each column. The final TMT sets were pooled from 14 single cells using a handheld multichannel pipette onto the prepared SOLAµ columns. After pooling, peptides were bound to the filter by centrifuging at 700g for 1 min. The SOLAµ plate was washed twice with 200 µL 0.1% FA, and the peptides were eluted twice with 20 µL of 40% ACN, 0.1% FA into an Eppendorf Twin.tec 96-well LoBind plate. Finally, the peptides were dried at 60°C in an Eppendorf Speedvac and reconstituted in A* for individual mass spectrometry analysis.

### CITE-seq sample preparation

Sorted Lin-CD34+ cells were washed with PBS and stained with Fc block for 10 minutes and then washed again. After Fc block the cells were stained for 30 minutes with the CITE-seq antibodies (**Table S2**). After the staining, cells were washed once again, counted to reach the proper concentration and loaded into the 10x instrument according to the manufacturer’s protocol. Single-cell RNA sequencing was performed using the 10x Genomics 3’ v3 platform. Approximately 7,000–10,000 cells were loaded in each 10x channel, after which the single-cell libraries were prepared according to manufacturer’s instructions, except for the added CITE-seq steps. A specific primer was added in the cDNA amplification step and instead of discarding the small fragments in the bead purification, they were kept and used for the CITE-seq/ADT library preparation according to the original publication^22^. After all libraries were ready, their quality and size were verified in the Bioanalyzer before sequencing on the NextSeq 500 Illumina platform. Two to three samples were pooled and sequenced together in each sequencing run.

### CRISPR/Cas9 targeting in human CD34+ BM cells

Normal BM was aspirated from a young, healthy donor and the CD34+ cells were isolated as described above. Cells were cultured for 2 days in StemSpan (Stem Cell Technologies, Cat. 09655) supplemented with TPO (100ng/mL), SCF (100ng/mL), FLT3 (100ng/mL), IL6 (100ng/mL) and UM171 (35nM) and electroporated with the ribonucleoprotein complexes (RNPs) made with 0.3 µl recombinant Cas9 (10µg/µl) using sgRNAs harboring 2’-O-Methyl at the first 3bps and last bp, 3’phosphorothioate bonds between the first 3bps and last 2bps as well as an 80-mer SpCas9 scaffold (Synthego). For the control locus AAVS1 we used a single sgRNA (5′-GGGGCCACUAGGGACAGGAU-3′), whereas we used pairs of sgRNAs for targeting H1F0 (5′-GAATTCCACGTCCGCCCCTGCGG-3′ and 5′-UUGGAGGCCUUGGCCCGCUUGGG -3′), SOD1 (5′-UAAUGGACCAGUGAAGGUGUGGG-3′ and 5′-CAUGAACAUGGAAUCCAUGCAGG -3′) and TALDO1 (5′-GCCGACACGGGCGACUUCCACGG-3′ and 5′-UCCAUCCUCUGACGCUUCACGGG-3′). Electroporation was carried out on an Amaxa D4 electroporator using the program DZ-100.

### Long-term culture initiating cell (LTC-IC) assay

One day post-electroporation, CD34+ cells were sorted (10, 30, 100, 300 and 1000 cells; 10 wells for each number) onto Mitomycin-C treated MS5 feeder cells in 96-well plates and cultured for 5 weeks in Gartner’s media (αMEM supplemented with 11% FCS, 12.5% Horse serum, 1% Pen/Strep, 1 uM hydrocortisone, 57,2 uM beta-mercaptoethanol, 20ng/mL TPO, 20ng/mL IL-3 and 20ng/mL GCSF) followed by 2 weeks in methylcellulose (MethoCult, Stem Cell Technologies, Cat. H4435). Stem cell frequencies were assessed by counting wells positive for cell growth.

### Colony Forming Cell (CFC) Assay

One day post-electroporation, 1,000 cells were plated in triplicate in 1 mL methylcellulose (MethoCult, Cat. H4435) and cultured for 2 weeks. Subsequently, different classes of colony-forming units (CFUs) were counted under a microscope.

### Mass spectrometry data acquisition

Peptides were loaded onto a 2cm C18 trap column (ThermoFisher 164946), connected in-line to a 15cm C18 reverse-phase analytical column (ThermoFisher EasySpray ES904), with 100% Buffer A (0.1% Formic acid in water) at 4 μl/min or 750 bar maximum pressure using the ThermoFisher EasyLC 1200, and the column oven operating at 30°C. Peptides were eluted over a linear 140-min gradient at a flowrate of 100 nL/min, using 80% Acetonitrile, 0.1% Formic acid (Buffer B) going from 10% to 30% B. Spectra were acquired with an Orbitrap Eclipse Tribrid mass spectrometer with FAIMS Pro interface (ThermoFisher Scientific) running Tune 3.4 and Xcalibur 4.3 using the RETICLE acquisition method^28^. FAIMS switched between compensation voltages of −50 V and −70 V with cycle times of 3 s and 1.5 s, respectively. MS1 spectra were acquired at 120,000 resolution with a scan range from 375 to 1500 m/z, normalized AGC target of 300%, and maximum injection time of 50ms. Precursors were filtered using monoisotopic peak determination set to peptide, charge state 2 to 6, dynamic exclusion of 120 s with ±10 ppm tolerance excluding isotopes and different charge states, and a precursor fit of 70% in a window of 0.7 m/z. Precursors selected for MS2 analysis were isolated in the quadrupole with a 0.7 m/z window. Ions were collected for a maximum injection time set to 46 ms and normalized AGC target of 300%. Fragmentation was performed with 30 normalized CID collision energy, and MS2 spectra were acquired in the linear ion trap (LIT) at scan rate rapid. LIT MS2 spectra were subjected to real-time search (RTS) using the homo sapiens database obtained from Uniprot (Swiss-Prot with isoforms, 42,297 sequences, downloaded on 30.03.2021), and trypsin set as enzyme (cleavage next to arginine or lysine, but not before proline). Static modifications were TMTpro16plex on Lysine (K) and N-Terminus and carbamidomethyl on cysteine (C). Oxidation of methionine (M) was set as variable modification. Maximum missed cleavages were set to 1 and maximum variable mods to 2. FDR filtering was enabled, maximum search time was set to 40 ms, and the scoring threshold was set to 1.4 XCorr, 0.1 dCn, and 10 ppm precursor tolerance. Use as trigger only was checked, and close-out was enabled with maximum number of peptides per protein set to 4. Precursors identified via RTS were isolated in the quadrupole with a 0.7 m/z window, ions were collected for a maximum injection time of 750 ms and normalized AGC target of 500%, fragmented with 32 normalized HCD collision energy, and MS2 spectra were acquired in the orbitrap at 120,000 resolution with first mass set to 120 m/z.

### Mass spectrometry data processing

Raw files were analyzed with Proteome Discoverer 3.1 (ThermoFisher Scientific) with the built-in TMTpro Reporter ion quantification workflows using the standard settings if not further specified. Reporter ion quantification was performed only on FTMS MS2 spectra, and the same spectra were also sent to Sequest for identification, where they were searched with precursor mass tolerance of 10 ppm and fragment mass tolerance of 0.02 Da. The database was the homo sapiens database obtained from Uniprot (Swiss-Prot with isoforms, 42,412 sequences, downloaded on 10.03.2024) with trypsin set as enzyme (cleavage next to arginine or lysine, but not before proline) and maximum number of missed cleavages set to 2. Static modifications were TMTpro16plex on lysine (K) and N-terminus, and carbamidomethyl on cysteine (C) was set. Dynamic modifications were set as oxidation (M) and Met-loss on protein N-termini. Results were rescored with Inferys^81^ and q-values were assigned with percolator and filtered to 1% FDR. For the reporter ion quantification, normalization mode and scaling mode were set to none and average reporter s/n threshold was set to 0. Isotopic error correction was applied.

### CITE-seq data processing

Demultiplexed FASTQ files were aligned to the human reference genome (GRCh38, v 2020-A as provided by 10x Genomics, which is based on GENCODE v 32/Ensembl 98) using Cell Ranger software (v 5.0.1) by 10x Genomics. The Cell Ranger “count” standard pipeline was used to obtain the expression matrix of unique molecular identifiers (UMIs) for each individual sample.

### scp-MS data analysis

scp-MS data was analyzed using the SCeptre pipeline^27^ using python 3.9 with SCeptre version 1.1.0, flowkit 0.9.3^82^ and Scanpy 1.9.1^29^. FACS .fcs files were processed in flowkit in the following manner. First, compensation and bi-exponential transform was applied and subsequently, data was merged and batch corrected using combat^83^ with each individual representing a batch. Subsequently, each FACS parameter in the merged data was transformed across single cells using the robust z-score with value clipping from -5 to 5. The resulting data was used to set the gates for the populations to annotate cells with a gated population. FACS data and sort- and label layouts were used to create the metadata for each cell.

Subsequent SCeptre analysis was performed in the following manner. A first pass of the SCeptre workflow was applied to identify clusters of cells that likely contain low quality cells (**Figure S1A**). Indications for low quality cell clusters were mixing of cells of unrelated FACS populations, low total protein signal and the combination of low FSC-A with high SSC-A. Cells contained in the low-quality clusters were filtered out in the second pass of the SCeptre workflow, which included the following steps. Raw files with less than 2,000 peptide-spectrum matches (PSMs) were removed and cells from low quality clusters were removed. To remove proteins whose values likely stem from contamination, a mean single cell to carrier ratio was calculated. For each protein in each cell, the signal ratio to the corresponding carrier channel was calculated. Proteins with a mean ratio of over 0.2 were removed. This also removed proteins where the ratio could not be calculated since there was no single cell signal in any cell. Subsequently, protein expression data was batch corrected using the SCeptre normalize function. Cells were filtered to have a signal for at least 600 proteins, and a minimum log2 total signal of 13.6 Furthermore, cells were filtered based on a relationship of FSC-A to log2 total signal (**Figure S1B**), as these parameters showed high correlation and thus were deemed to be suitable to filter outlier cells. The relationship between the two parameters was modeled using the function sklearn.linear_model.RANSACRegressor and cells with a prediction error of over 0.8 log2 total signal were removed. Finally, proteins were required to have a signal in at least 10 cells. This resulted in a dataset of 2,934 proteins across 2506 cells with various data completeness (**Figure S1C**) and for the second dataset 2174 proteins across 922 cells.

Subsequently, protein expression data was median-ratio normalized and log2(x+1) transformed. This represents the “batchcorr_norm_log2” layer that was used for differential expression testing. For the embedding, data was imputed using the k-nearest neighbor algorithm followed by the application of the “regress_out” function of Scanpy using log2 total signal and number of proteins with signal as parameters. To embed the data, principal component analysis (PCA) was applied which was used to calculate a neighborhood graph, which was subsequently embedded using UMAP. Unsupervised cell clustering was performed using the Leiden algorithm. Clusters were manually annotated using the expression of marker genes and FACS information.

The quality of batch effect removal was quantified as the overall variance contribution given a covariate (PC regression value) as implemented in the python single-cell integration benchmarking package scib^31^.

Differential expressed proteins in each cluster were calculated by comparing protein expression in each cluster against all other clusters using the layer “batchcorr_norm_log2”. For each protein, a two-sided Welch’s t-test was performed, excluding missing values. P-values were corrected via the Benjamini–Hochberg procedure and a cutoff of 5% FDR was applied.

To compare protein log2 fold-changes derived from scp-MS to bulk data, proteomics data^42^ from bulk sorted human bone marrow populations was used, which was reanalyzed with Spectronaut 18.4 using factory settings. Replicates were aggregated to the mean and the log2 fold-change of each protein in each population was calculated by subtracting the mean of all other populations. scp-MS data was matched to the smallest matching bulk protein group. For each scp-MS cluster and gated population log2 fold-changes were compared for proteins that had at least 30% coverage in the scp-MS cluster and were detected in the bulk dataset. Each scp-MS cluster and gated population was compared to each bulk population by calculating the Pearson correlation and the resulting matrix was clustered hierarchically.

For the unsupervised clustering of proteins, proteins were filtered to have at least one significant log2 fold-change above 0.2 (30% cluster coverage, 1% FDR) in any of the cell clusters. For each cell cluster, values from layer “batchcorr_norm_log2” were aggregated to the mean ignoring missing values and each protein was z-score transformed across cell clusters. Proteins were clustered hierarchically using Euclidean distance as metric and the linkage method ward and the tree was cut to 9 protein clusters. Each protein cluster was tested for enriched GO Biological Process and KEGG terms using gseapy^50^. The background was set to all proteins in the dataset and terms were filtered 5% FDR and more than one term per gene set. For each protein cluster selected terms were plotted.

### CITE-seq data analysis

The CITE-seq expression matrices were analyzed using scanpy 1.9.8^29^ and scvi-tools 1.1.2^55^. Cells were filtered to contain at least 1,800 mRNA counts, at least 800 genes and a fraction of mitochondrial gene counts between 0.025 and 0.1. Genes were filtered to be detected in at least 99 cells. Cells were annotated via Azimuth^4^, separately for each individual. We removed cells that were annotated as a rare celltype or celltypes with no evidence in the scp-MS data. Only CITE-seq surface markers that overlapped with scp-MS markers were retained. Genes were mapped to the scp-MS data using an ENSG-Uniprot mapping table. If multiple proteins or mRNAs mapped to the same ENSG, the protein or mRNA with the highest coverage was mapped. Subsequently, we created the “counts_totalnorm_log2” layer by normalizing each cell to total counts and applying log(x+1) transform. Next, genes were subsetted to contain 4,000 highly variable genes with flavor “seurat_v3” and batch keys provided, and to contain all genes detected in scp-MS, resulting in 5820 genes. Count data was modeled and batch-corrected using totalVI^52^. The modeled mRNA expression was median-ratio normalized and log2 transformed, resulting in the “denoised_rna_norm_log2” layer. For the UMAP embedding, the modeled mRNA expression was used. Azimuth GMPs were subsetted into GMDP and MDP using leiden clustering. Cell cycle phase was predicted using a list of marker genes and the “score_genes_cell_cycle” function of Scanpy. To benchmark mRNA log2 fold-changes from either counts or scVI-modeled data, transcriptomics data^42^ from bulk sorted human bone marrow populations was used. Transcriptomics data was CPM normalized and log(x+1) transformed. Log2 fold-change between two populations was calculated by subtracting the means of the replicates.

### Integration of scRNA-seq and scp-MS

Integration was performed using GLUE^56^. The guidance graph was constructed to contain all 2,934 proteins of scp-MS and all 5,820 mRNAs from scRNA-seq (CITE-seq), resulting in 8,754 nodes. 2,778 edges with weight 0.9 were inserted between protein and matching mRNA. Self-loop edges with weight 1.0 were inserted for each node. This resulted in 11,532 edges. Further required input for GLUE were an expression matrix and a latent representation thereof per modality. For scp-MS, the imputed data that was not processed with “regress_out” was used as expression matrix and as latent representation, 50 PCs of that were used. The probabilistic generative model used by the decoder was set to “Normal” corresponding to a “LogNormal” distribution considering the prior log-transformation of the scp-MS data. For scRNA-seq, the count data was used as expression matrix and the totalVI latent space was used as representation, which provided additional batch correction. The probabilistic generative model was set to negative binominal (“NB”) and the individuals were defined as batch variable. In contrast to the standard GLUE workflow that used a two-stage training procedure to estimate balancing weights, we used a single-stage procedure without balancing weights. The GLUE integration resulted in a joint latent representation of “protein” and “mRNA” cells. After integration, scRNA-seq labels were transferred to scp-MS using the GLUE function “transfer_labels” using the GLUE latent space and 15 nearest neighbors. Modality integration as well as biological signal preservation were measured using scIB metrics^31^. To quantify the overlap of the distribution of cells from the two modalities in the integrated latent space, we used the average silhouette width score that compares the average distance between cells of the same modality to the average distance of cells from different modalities. The highest possible score for separation of the modalities is 1, while values near 0 indicate well overlapping sets of cells. To assess preservation of biological variance, we compared cell type separation metrics before and after integration per modality. For the “mRNA” cells, we used the Azimuth-derived cell type annotations as ground truth that are to be preserved and the totalVI embedding as baseline. For the “protein” cells, we used the leiden cluster-based cell types and the PCA embedding that was also used as input to GLUE. We computed the average silhouette width score as just described as well as the normalized mutual information and adjusted rand index scores as implemented in scIB. Both compare the overlap of two clusterings. One being the ground truth cell type annotations and the other one being derived from the embedding structure as data-driven clusterings. A perfect match of the clusterings would lead to a score of 1, uncorrelated or random clusterings to a score of 0.

### Cellrank analysis

A cellrank^5,6^ analysis was performed on the integrated dataset, with standard setting applied if not further specified. First, the joint latent space was used to calculate a nearest neighbor graph, following a diffusion map. Then, after setting an immature HSC as root cell, a diffusion pseudotime was calculated, which was used to initialize a cellrank pseudotime kernel. Subsequently, a transition matrix was computed and macrostates were determined via GPCCA. Terminal states were determined by fitting seven macrostates to a Schur decomposition and selecting the top six. Finally, fate probabilities were calculated for the six terminal states.

Expression profiles were modeled using the approach in cellrank where a generalized additive model (GAM) is fitted to the expression values along pseudotime, while using the fate probabilities as weights. Cells were subsetted into the relevant clusters for each trajectory. GAM settings were 12 knots, lam=12 and derivative was set as penalty. For mRNA, the scVI-modeled values were used. For protein, missing values were ignored by setting the weights of cells with missing values for the specific protein to zero. For both modalities, values were not log2 transformed, but used in linear space. Furthermore, values were normalized by dividing them by the global mean of the specific gene.

HSC quiescence and differentiation was investigated using a subset of cells with pseudotime below 0.4. Each protein or mRNA had to be detected in at least 200 cells to be considered. Next, the correlation vector was calculated by calculating the Pearson correlation of each log2 transformed mRNA/protein with pseudotime. For protein, missing values were ignored for this calculation, while for the count-based mRNA, zeros were used. scVI modeled mRNA did not have any missing values.

Lineage specification was investigated using all cells, however using only mRNA/proteins with detections in at least 200 cells. Correlations vectors for each lineage were calculated by calculating the Pearson correlation of each log2 transformed mRNA/protein with fate probability. For protein, missing values were ignored, while for the count-based mRNA, zeros were used. scVI modeled mRNA did not have any missing values.

Gene set enrichment analysis (GSEA) was performed on the correlation vectors using GSEApy^50^. A preranked GSEA was computed using the gene set GO Biological Process with minimum set size of 3 and maximum of 1000. Resulting terms were filtered to 10% FDR and for the lineage specification additionally to a positive enrichment score. The term list was shortened by removing redundant GO term using REVIGO^84^.

Gene expression cascades for each trajectory were visualized as heatmaps by filtering the correlation vector to a minimum correlation of 0.25 for mRNA and 0.2 for protein. Then genes were sorted by the pseudotime coordinate of their maximum expression in the modeled expression profile. Then, this vector was binned into 4 bins and the top 5 mRNAs or top 8 proteins of each bin were selected for visualization.

For the joint mRNA-protein covariation analysis the protein lineage specification correlation vectors were concatenated into a table of protein lineage correlations across the six lineages. In each vector column, if the protein did not pass the 200-cell coverage filter, it was set to missing. The table was subsequently filtered for proteins with Pearson correlation of at least 0.2 in any of the six lineages. For mRNA the count-based correlation vectors were used. Then the intersection of both protein and mRNA tables was concatenated and a pairwise correlation matrix was computed, while requiring at least four values to return a Pearson value. Keeping only columns with proteins yielded a correlation vector of a protein with all other mRNAs and proteins and sorting this vector by correlation resulted in a correlation rank for each mRNA and protein. For each protein, related proteins were defined to be in the same KEGG pathway.

To compare the mRNA correlation ranks for each protein to results from bulk data, we used a dataset containing mRNA and protein measurements of 59 breast cancer cell lines^77–79^. Log2 transformed tables were normalized by subtracting the row and column medians. Similar to the single-cell data, tables were concatenated, and a correlation matrix was computed and an mRNA correlation rank for each protein was computed. Ranks were then compared between bulk and single-cell data.

### Gene expression modelling with scProtVelo

#### Creating paired single cell data

To create a dataset with paired mRNA and protein expression for all cells, we selected cells of the erythroid trajectory based on a leiden clustering of the GLUE embedding, so that we obtain the same biological subset of cells from both modalities. This cell subset contained 483 cells from the protein dataset and 1,729 cells from the scRNA-seq data. We selected 1,068 proteins that were detectable in at least 30% of the protein cells within this subset and 5,117 mRNAs that had non-zero counts in at least 2% of the scRNA-seq cells, which gave us a total of 1,039 overlapping genes to work with. mRNA counts and batch corrected proteins were median-ratio normalized and, as also done for RNA velocity, expression values were smoothed per modality over their respective nearest neighborhood graphs. Then, for each cell, values for the not measured modality were interpolated over the 15 nearest neighbor cells of the modality of interest based on the joint GLUE embedding. For all features, 25% and 75% quantiles were computed and used to scale features by 1/(75% *quantile* − 25% *quantile*) to give all features the same weight in the scProtVelo reconstruction loss later on. We want to note here that scaling of expression values only results in scaled transcription and translation rates that can be reverted after model fitting, while a shift cannot be compensated for within the differential equations we define below.

We further define genes to showcase significant variation during the selected differentiation process, if they are differentially expressed between the upper and lower 30% quantiles of cell with respect to diffusion pseudotime in both modalities. For that we randomly sampled 50 cells from the respective quantiles and selected genes at a significance level of 5% after Benjamini-Hochberg multiple testing correction that had a log-fold-change of at least 0.1 on protein level and 0.5 on mRNA level. This resulted in a selection of 147 genes.

#### The translation dynamics model

In a first modelling setup, we followed the concept of RNA velocity^7^. We jointly model mRNA *r*(*t*) and protein *p*(*t*) expression of a gene considering the transcription of a gene to mRNA molecules at rate *α*, their degradation at rate *β* as well as translation to proteins at rate *κ* and their respective degradation at rate *δ* over time t. We assume fixed (time independent) rates per gene and also model time as gene specific, since orthogonal processes like cell cycle as well as cell differentiation could be the driving factors in different genes. Per gene, this process can be formalised by the following ordinary differential equations:

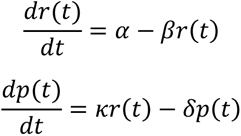

The expression values *r*(*t*) and *p*(*t*) stand for nearest-neighbour smoothed median-ratio normalised abundances. Both, the time annotations t as well as the kinetic rates *α*, *β*, *κ* and *δ* are not directly measured and are to be inferred.

Given that we model fixed, time-independent rates, a gene can either be in an induction or repression state, characterised by mRNA expression increasing or decreasing with time. In the RNA velocity formulation, induction is defined to start with unspliced as well as spliced expression being 0. We observe different behaviour in mRNA-protein phase portraits. Thus, we assume free initial expression values *r*_0_*^ind^* and *p*_0_*^ind^* and pose the restriction on the kinetic parameters that the derivatives 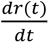 and 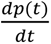 are non-negative at *r*_0_*^ind^* and *p*_0_*^ind^*, respectively. For the case of repression, we model the initial expression values *r*_0_*^rep^* and *p*_0_*^rep^* as a result of an unobserved induction phase that shares all kinetic parameters, but the transcription rate, with the following repression phase and enforce that the derivative of the mRNA expression is non-positive at the inferred starting point (*r*_0_*^rep^*, *p*_0_*^rep^*). Additionally, similar to the RNA velocity model, we explicitly also allow cells to be in a steady state that we set to be the respective *r*_0_ and *p*_0_ values depending on the general gene state.

On top of this very free model setup, we used a second model variation that is meant to capture one known linear process more accurately. In this setting, we restrict time to be shared across all genes and introduce diffusion pseudotime as a prior on the modeled time.

#### scProtVelo

We implemented all those considerations into scProtVelo. The scProtVelo model, in short, is a variational autoencoder-like architecture that takes a cell representation as input, learns a latent representation of it that is further used to generate mRNA and protein expression that is close to the original expression of that cell. The crucial part of scProtVelo is that the generation is not performed by a fully connected neural network. Instead, it infers a latent time annotation for a cell as well as the kinetic parameters and combines both to generate the expression values based on the defined expression dynamics. Our python implementation heavily builds on the veloVI framework, and we refer the reader to the original publication^9^ for all technical modelling details that are not explained in the following.

#### The scProtVelo generative model

The differential equations introduced above can be solved analytically and as

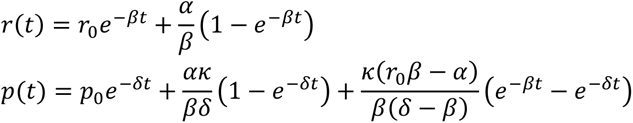

with initial expression values *r*_0_ and *p*_0_and as such build the base for the generative process of scProtVelo. We can now formalise all four possible cell-gene states (induction steady state, induction dynamic state, repression steady state, repression dynamic state) described by the gene specific variable *k^ind/rep^* and the gene and cell specific variable *k*_*i*_^*ss/ds*^for a cell *i* ∈ *I*. Time *t* as well as kinetic rates *α*, *β*, *κ* and *δ* and intermediate variables are gene specific, but we omit the subscripts for readability. For the induction steady state, we propose a restriction-free expression pair

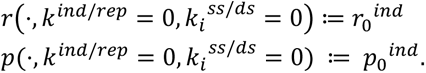

For the corresponding dynamic case, mRNA and protein expression can be described to follow the analytical solution given above with the initial state being the values defined for the steady state case:

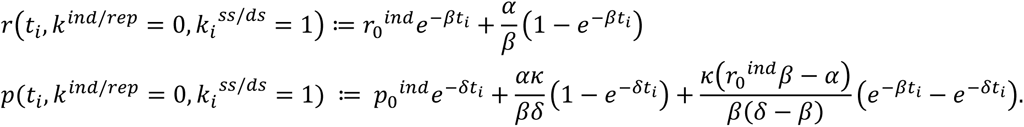

For the repression modelling, we assume an unobserved induction phase preceding the observed repression phase. We assume that the induction phase shares rates *β*, *κ* and *δ* with the repression phase and that both phases just differ in their transcription rates *α_ind_* and *α_rep_*. Further assuming an initial state 0, 0 for the induction and a gene-specific time 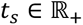 at which the initial expression pair is attained, we propose the repression steady state to be

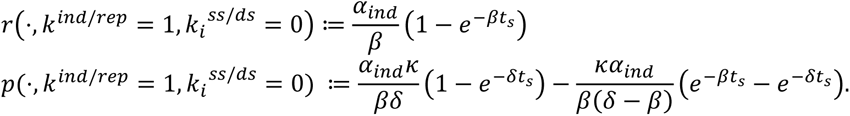

Last, for the dynamic repression state, we obtain

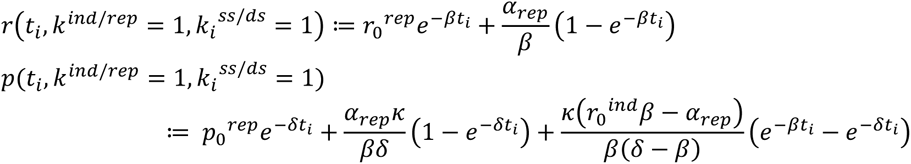

with *r*_0_*^rep^* and *p*_0_*^rep^* being the values obtained above for the repression steady state.

For the generative process of an expression pair (*r_i_*, *p_i_*) of a cell *i*, we draw a low-dimensional (*d* = 10) latent cell representation

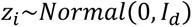

as well as a cell-independent distribution over gene states (induction or repression) described by the probability simplex generated by the Dirichlet distribution and then the actual state assignment itself

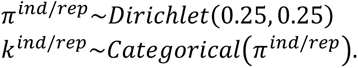

Further, we draw a distribution distinguishing between a cell being in steady state or dynamic state as well as the assignment

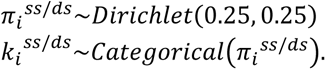

The time annotation *t_i_*, we model as a function of the latent cell representation *z_i_*

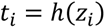

with 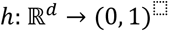 being a fully connected neural network.

Finally, the expression pair (*r*_*i*_, *p*_*i*_ ) is drawn from two normal distributions as

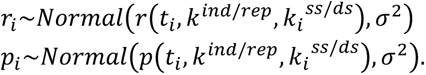

#### scProtVelo inference

The goal of the inference is to obtain estimates for all introduced kinetic rates, gene state distribution parameters, neural network weights, initial expression values for the induction cases, switch times for the repression cases and finally posterior distributions over the latent variables *z*, *π*^*ind/rep*^ and *π^ss/ds^*. We use variational inference and with *x_i,glue_* being a precomputed representation of the mRNA and protein expression of cell *i* ∈ *I*, namely its GLUE representation, we propose the following factorization of the posterior distribution

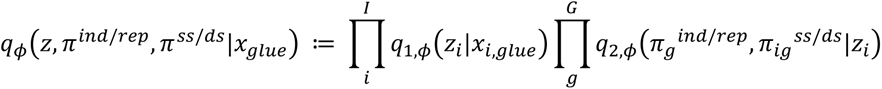

meaning that a neural network *q*_1_ maps a precomputed cell representation *x_i,glue_* to a posterior distribution for the latent cell representation *z_i_* which is itself used as input to a second neural network *q_2_* that maps it to posterior distributions for the state distributions *π*_*g*_^*ind/rep*^and *π*_*ig*_^*ss/ds*^ for all genes *g* ∈ *G*. For the likelihoods we integrate over the four defined cell-gene states and obtain:

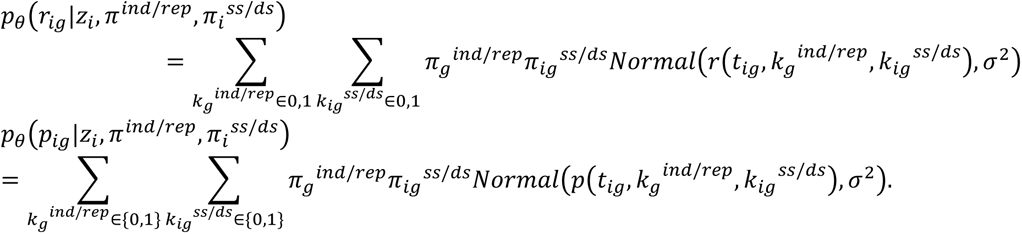

Additionally, to minimizing the arising ELBO objective that is composed of the negative log likelihood of the data given the fitted model as well as Kullback-Leibler divergence terms for the latent variables *z*, *π^ind^*^/*rep*^ and *π*^ss/ds^, we include two more loss terms to be optimised. Firstly, we add *L_param,ind_* to enforce non-negative derivatives in mRNA and protein expression at the initial values in the induction case and *L_param,rep_* to ensure non-positive derivatives in mRNA expression at the initial values in the repression case

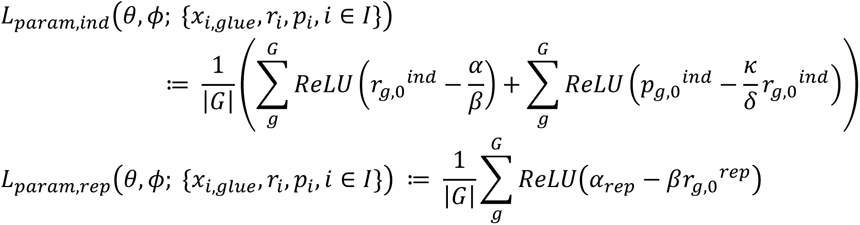

with loss values set to 0 whenever a denominator is 0 and weighted strongly enough to enforce all wanted conditions.

Secondly, we introduce *L_init,ind_* and *L_init,rep_* as penalty terms used during a pretraining phase that encourages the modeled steady states to coincide with heuristically observed steady state values *r_g_^high^*, *r_g_^low^*, *p_g_^high^* and *p_g_^low^* that are computed as the observed median expression of the cells that have mRNA expression within the highest or lowest 1% quantile of that gene

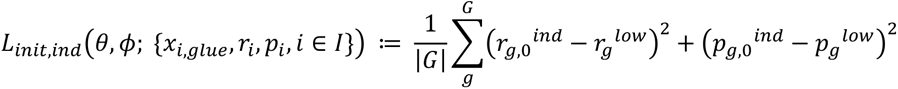

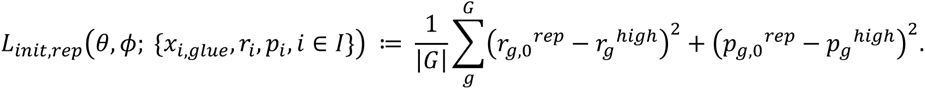

#### Trajectory inference using scProtVelo

In the trajectory inference setting, we start model fitting with a pretraining phase with a high focus on the *L_init_* loss terms, because we noticed that it helped general model fitting to get initial states approximately correct first. During normal training, this loss term is then turned off.

To quantify the performance in inferring the correct direction of cell progression, we subsetted all genes we fitted the model on to the ones that we accepted as changing along the differentiation trajectory and computed the direction of change as the sign of the difference between mean expression within the late 20% quantile with respect to pseudotime and the mean within the early 20% quantile of the cells. For the predicted state, we chose the state that had the higher value in *π*_g_^ind/rep^. To get an impression of the model performance, we sorted the genes by their model likelihood which represents a measure of how well the model captured the expression values of a gene, without knowing whether the direction of change was inferred correctly or not. Finally, we computed accuracy scores on increasing numbers of genes and compared it to the expected accuracy of a random permutation of the model predictions.

We based velocity computations on the 100 genes with the highest likelihood values and followed the same procedure as done for RNA velocity in the veloVI framework.

For the comparison to RNA velocity results, we followed the proposed veloVI workflow on unspliced and spliced mRNA counts of the mRNA cells that were used in the scProtVelo run. To create the .loom files, velocyto 0.17.17^7^ was applied on the cellranger count output. To compare the accuracy of the cell-to-cell transition graphs that the stream plot visualizations are based on, we checked whether proposed transitions from one cell to a neighboring cell coincide with an increase in diffusion pseudotime or not. We aggregated these values to accuracies per cell that we inspected visually projected on the respective UMAP embeddings as well as compared their distributions in a histogram.

#### Gene expression modelling with scProtVelo

To quantify the variance in protein expression that is determined by the corresponding mRNA expression, we figured it made only sense to consider genes that exhibit biological variation and restricted our analysis on the genes we found to be differentially expressed between the start and the end of the chosen differentiation trajectory. To ensure further that we quantify the variance corresponding to cell differentiation, we restricted latent time inference to only one shared time shared across all genes guided by a diffusion pseudotime prior.

To accommodate for those model specification changes, we firstly included a loss term representing the diffusion pseudotime prior t_*ii*_^dpt^, i ∈ *I* as

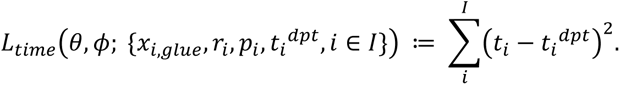

Also, the shared time stands in conflict with the prior implementation of the steady states. So instead of modelling a steady state vs dynamic state variable per cell and gene, we learn a switch time per gene at which gene expression for a gene switches from a constant steady state value to dynamic expression. For induction state, this switch time is restricted to be within (0, 1) just as the fitted time. For the repression case, we model a switch time in (−∞, 1], where a negative value implies an unobserved earlier initial state and observed cells just cover a more advanced time range. This is essential considering that the initial state for repression is tied to an induction phase. And finally, having access to diffusion pseudotime, we also refine on the heuristically estimated initial states and use the median expression of the cells within the 1% quantile with respect to diffusion pseudotime.

During training we leave out 20% of the cells at random as a test set. To compare protein estimates from scProtVelo against a linear model, we select the protein cells to avoid the computationally interpolated protein values. We then log transform mRNA and protein expression values and fit a linear model (including a bias term) on the train set. Finally, we compute R^2 values for protein predictions from both models in log expression space.

---

**Figure S1.**
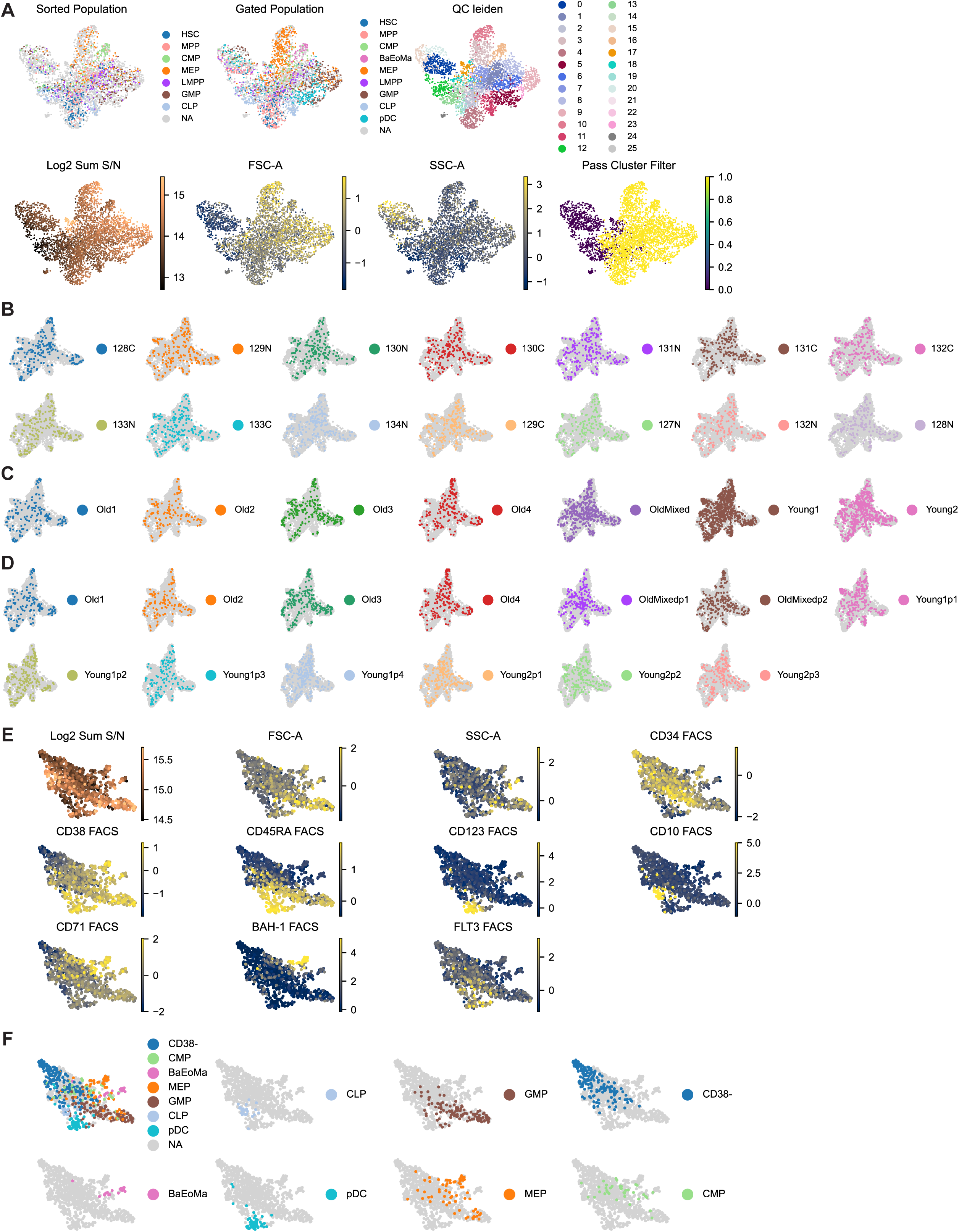
SCeptre processing of scp-MS dataset and analysis of second scp-MS dataset with new FACS markers, related to Figure 1. (A) UMAP of cells from the first pass of the SCeptre workflow. Cell clustering is used to remove spurious clusters of low-quality wells/cells. (B-D) UMAP of cells annotated with their respective TMTpro channel, individual donor (C) and 384-well plate (D). OldMixed was sorted from a pool of cells from Old1-4. (E) UMAP of cells of the second scp-MS dataset with new FACS markers, annotated with total signal and FACS parameters. (F) UMAP annotated with the gated populations.

**Figure S2.**
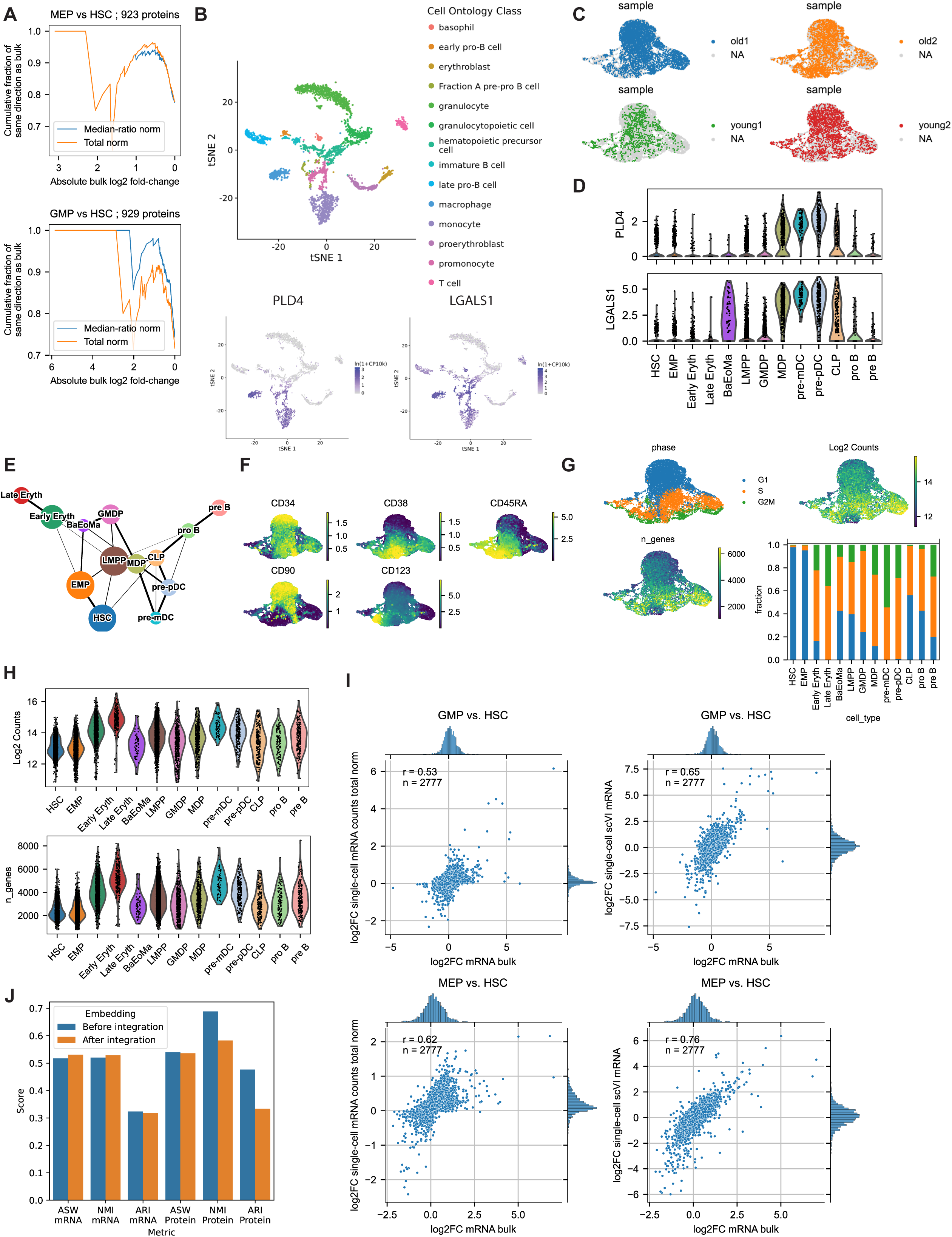
Additional analyses related to Figure 2 and Figure 3. (A) Benchmark of cell expression normalization in scp-MS by normalizing to total signal or median-ratio. We compared the fold-change direction between the gated single-cell populations to sorted bulk data and found median-ratio normalization to yield more proteins which change points into the same direction as in bulk data. (B) Branching of MDPs from GMDPs is characterized by the expression of LGALS1 and PLD4 as observed in the Tabula Muris scRNA-seq dataset: tabula-muris.ds.czbiohub.org. (C) Donor distribution across UMAP embedding, related to Figure 3. (D) Separation of MDPs from GMDPs by the expression of PLD4 and LGALS1 as in the scp-MS dataset. (E) PAGA graph of celltype labels. (F) totalVI modeled CITE-seq surface marker expression. (G) UMAP with overlaid cell cycle phase, total counts per cell, number of detected genes per cell, and barchart with cell cycle phase distribution. (H) Total counts and genes per population. (I) mRNA fold-changes between populations based on single-cell data were compared to CPM normalized bulk data^39^. Left column represents single-cell data based on log2(x+1) counts data normalized to total counts. Right column represents scVI-modeled data in log2(x) and median-ratio normalized. Top row represents the comparison of GMP to HSC, with GMDP selected as single-cell population. Bottom row represents the comparison of MEP to HSC, with Early Eryth selected as single-cell population. (J) Metric on celltype separation before and after GLUE integration for CITE-seq and scp-MS labels. ASW=Average silhouette width, NMI=Normalized mutual information, ARI=Adjusted rand index.

**Figure S3.**
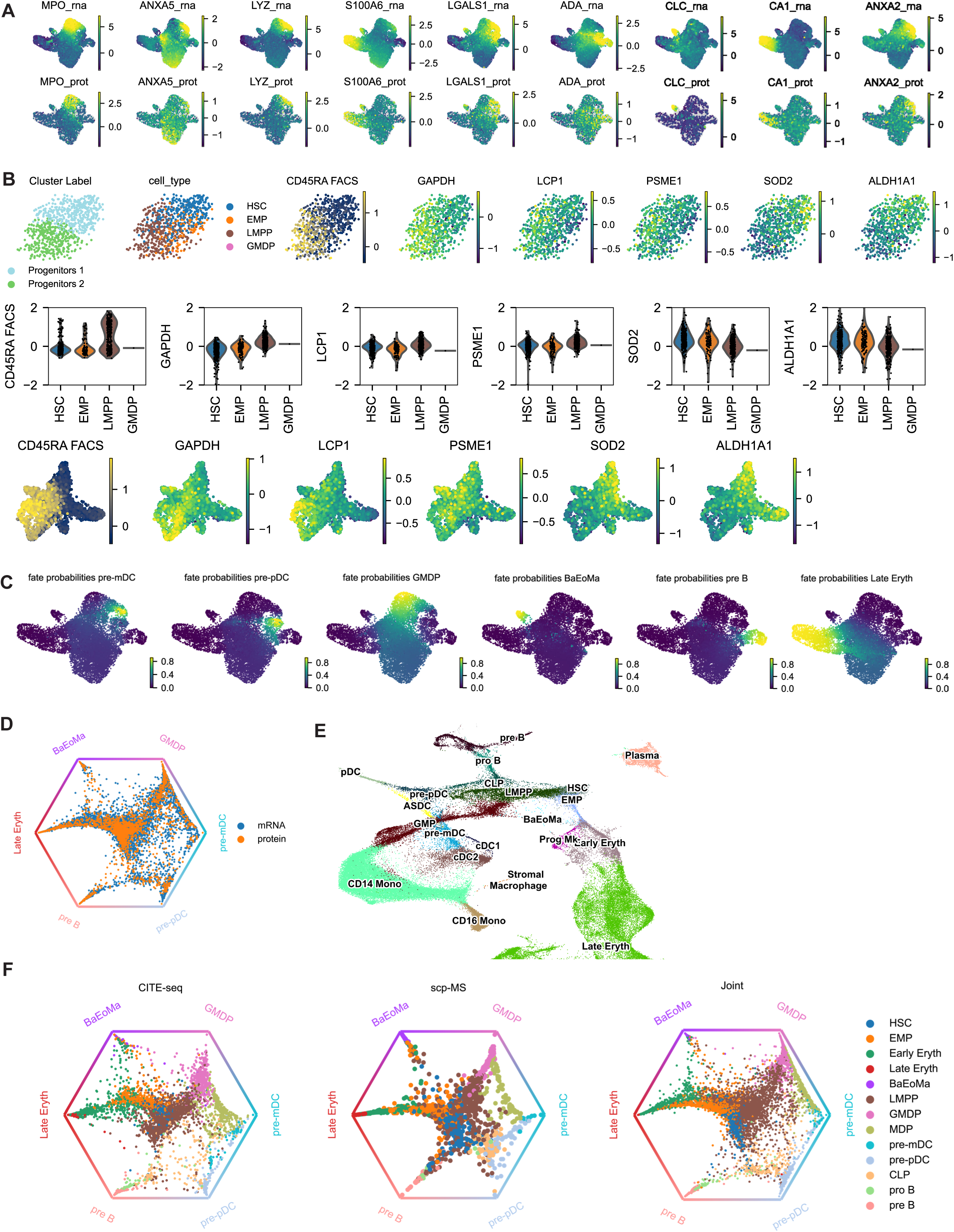
Additional analyses related to Figure 3 and Figure 4. (A) UMAP of selected protein markers that correspond well to mRNA expression. (B) Validation of cell label transfer of scRNA-seq labels EMP and LMPP onto scp-MS labels Progenitors 1 and 2, related to Figure 3. (C) Visualization of fate probabilities, related to Figure 4. (D) Cellrank circular projection with overlaid “mRNA” and “protein” cell labels. (E) UMAP representation of Azimuth map. (F) Comparison of fate probabilities of different cellrank models. Terminal states determined in the joint model were imposed on the CITE-seq and scp-MS cellrank model and fate probabilities into those terminal states were calculated. The circular projection was used to compare the fate probabilities, which revealed the better separation of pre-mDC and pre-pDC terminal states in scp-MS and joint data compared to CITE-seq.

**Figure S4.**
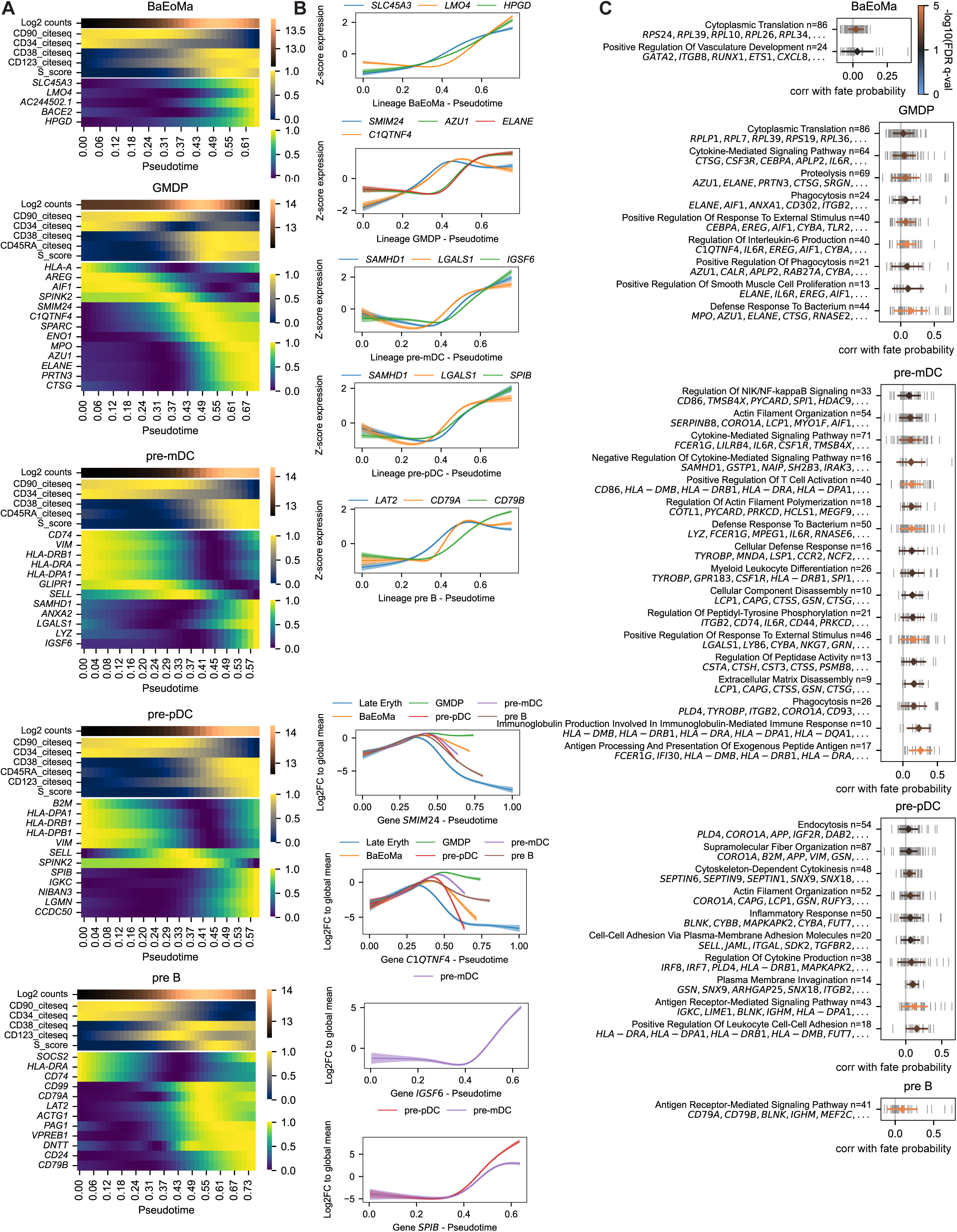
mRNA level analysis of cellrank trajectories on joint latent space, related to Figure 4. (A) Left column shows heatmap of mRNA expression cascade for each lineage. (B) Middle column shows the modeled mRNA expression profiles for each lineage and expression profiles of a gene across all lineages. (C) Right column shows GSEA results of each lineage.

**Figure S5.**
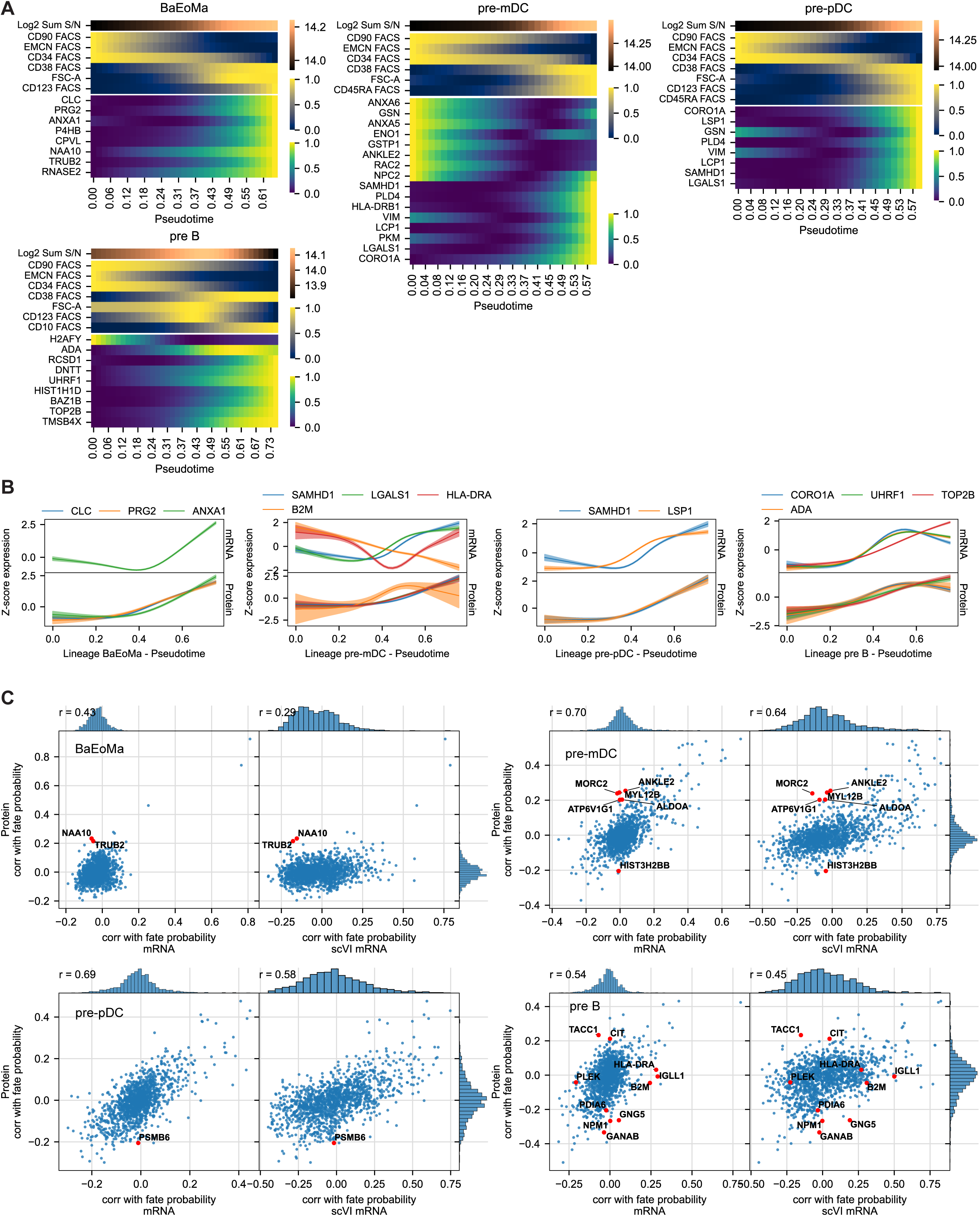
Protein level analysis of cellrank trajectories on joint latent space, related to Figure 6. (A) Heatmaps of protein expression cascade. (B) Modeled protein expression profiles of selected proteins. (C) Scatterplot comparing the correlation vectors between mRNA and protein level.

**Figure S6.**
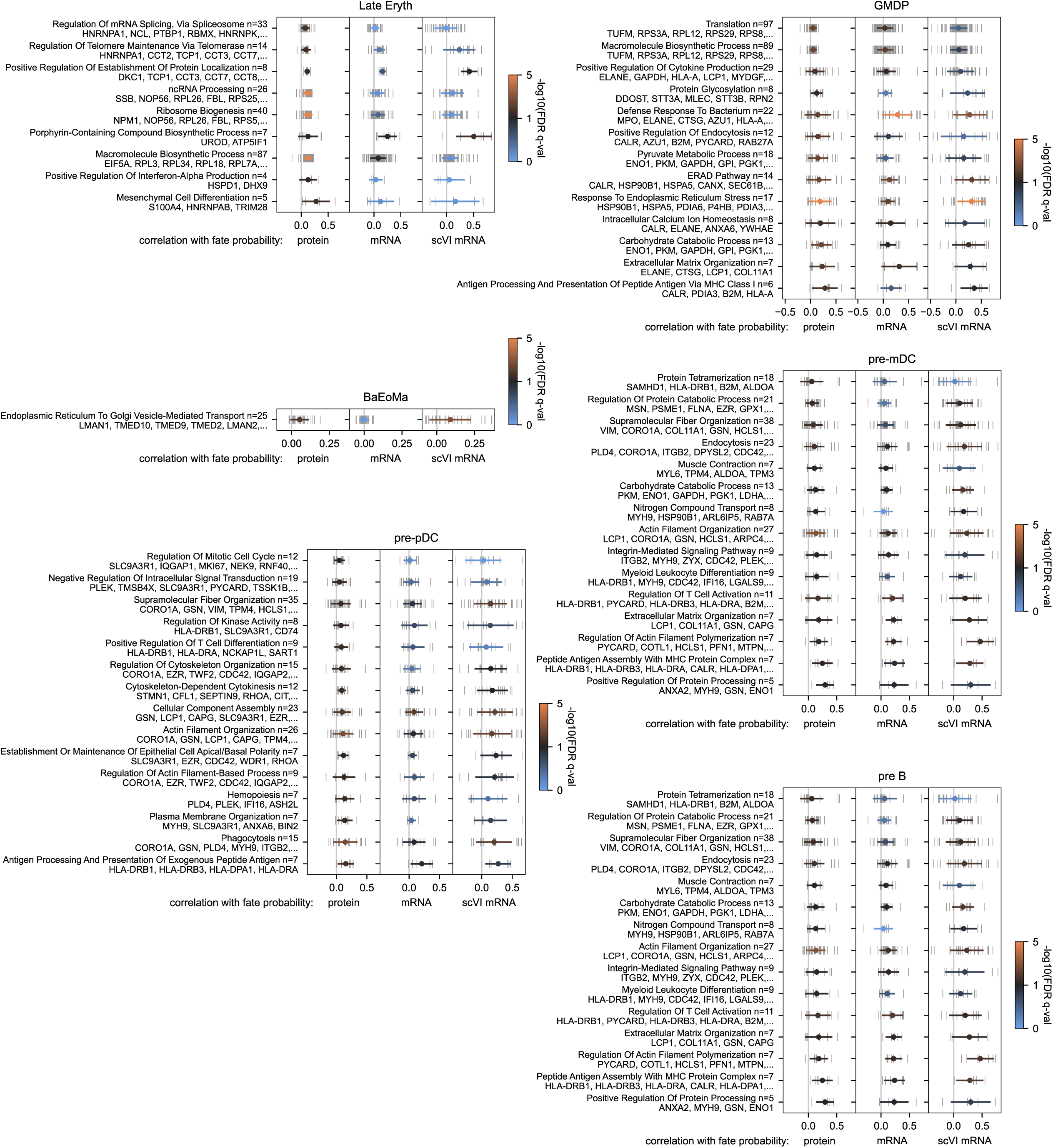
Further protein level analysis of cellrank trajectories on joint latent space, related to Figure 6. GSEA results for the lineages compared between mRNA and protein.

**Figure S7.**
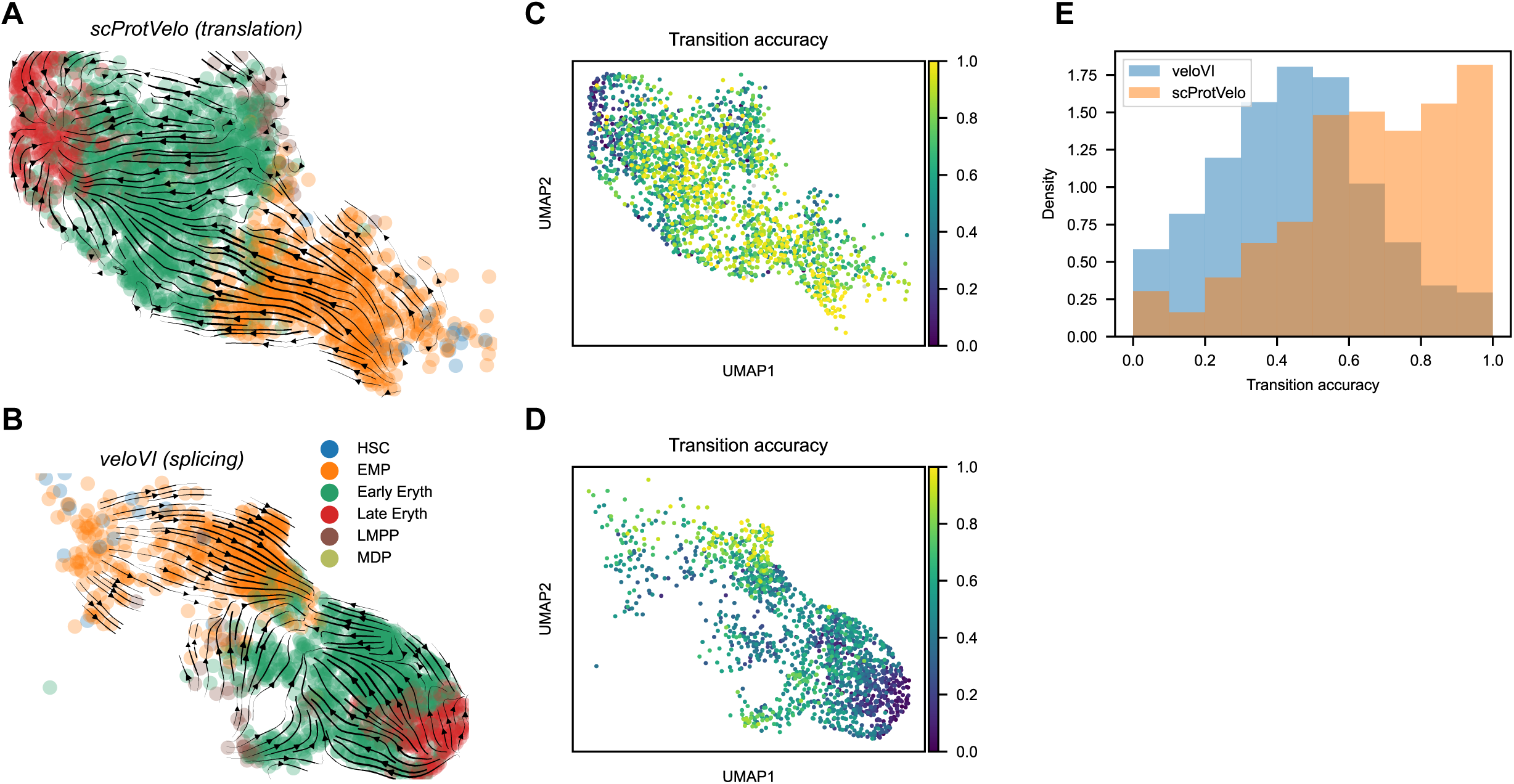
Further quantitative comparison of trajectory inference results from scProtVelo and veloVI, related to Figure 7. (A, B) Translation-based (A) and splicing-based (B) velocities projected onto a UMAP embedding. (C, D) Cell transition accuracies based on scProtVelo (C) and veloVI (D) projected onto a UMAP embedding. (E) Density histogram of the accuracy values from both approaches.

